# Genome-wide Detection of Cytosine Methylations in Plant from Nanopore sequencing data using Deep Learning

**DOI:** 10.1101/2021.02.07.430077

**Authors:** Peng Ni, Neng Huang, Fan Nie, Jun Zhang, Zhi Zhang, Bo Wu, Lu Bai, Wende Liu, Chuan-Le Xiao, Feng Luo, Jianxin Wang

## Abstract

Methylation states of DNA bases can be detected from native Nanopore reads directly. At present, there are many computational methods that can detect 5mCs in CpG contexts accurately by Nanopore sequencing. However, there is currently a lack of methods to detect 5mCs in non-CpG contexts. In this study, we propose a computational pipeline which can detect 5mC sites in both CpG and non-CpG contexts of plant genomes by using Nanopore sequencing. And we sequenced two model plants Arabidopsis thaliana (*A. thaliana*) and Oryza sativa (*O. sativa*) by using Nanopore sequencing and bisulfite sequencing. The results of our proposed pipeline in the two plants achieved high correlations with bisulfite sequencing: above 0.98, 0.96, 0.85 for CpG, CHG, and CHH (H indicates A, C or T) motif, respectively. Our proposed pipeline also achieved high performance on Brassica nigra (*B. nigra*). Experiments also showed that our proposed pipeline can achieve high performance even with low coverage of reads. Moreover, by using Nanopore sequencing, our proposed pipeline is capable of profiling methylation of more cytosines than bisulfite sequencing.

## Introduction

As a major type of DNA methylation, 5-methylcytosine (5mC) plays important roles in biological processes of animals and plants^1,2^, such as the regulation of gene expression^3^ and silencing of transposable elements^4^. In animals, 5mC primarily occurs at CpG sequence contexts^5-7^. In contrast with animals, both CpG and non-CpG (CHG and CHH, where H is A, C or T) sites can be significantly methylated in plants^8,9^. It is reported that there exists widespread variation in DNA 5mC between plant species^10^. For example, there are overall genome-wide levels of 24% CpG, 6.7% CHG and 1.7% CHH methylation in Arabidopsis thaliana (*A. thaliana*)^11^, while Beta vulgaris (*B. vulgaris*) has 92.5% CpG, 81.2% CHG and 18.8% CHH methylation level^10^. The establishment and maintenance of non-CpG methylation is due to chromomethylases and RNA-directed DNA Methylation (RdDM) pathways^12^. Non-CpG methylation regulates both transposable elements and genes^4^, which can ensure genome integrity and influence reproduction and development of plants^12,13^. It is also reported that, compared to CpG methylation, non-CpG methylation can silence transposable elements stronger and is crucial for silencing CpG-depleted transposable elements^14^.

Bisulfite sequencing is currently the gold standard technique for profiling methylation of cytosines^15^. In a bisulfite treated genomic DNA, unmethylated cytosines are converted to uracils, while methylated cytosines are unchanged^16^. 5mC in both CpG and non-CpG contexts can be detected by bisulfite sequencing. However, there are also drawbacks of bisulfite sequencing, such as short reads, incomplete conversion, and DNA degradation during bisulfite treatment, which leads to lack of specificity and loss of sequencing diversity^17,18^. Recently, Nanopore sequencing from Oxford Nanopore Technologies (ONT) offers great opportunities for DNA methylation detection^19^. The chemical modification to DNA can affect the electrical current signals near modified bases in Nanopore sequencing^20^. Thus, DNA methylation can be directly detected from native DNA reads of Nanopore sequencing without extra laboratory technique, which makes DNA degradation and amplification biases be avoided. Moreover, the long reads of Nanopore sequencing makes it possible to profile methylation in repetitive or low complexity regions^21^.

Methods using Nanopore sequencing for DNA 5mC detection can be classified into three categories: (1) Statistics-based methods, such as Tombo^22^, infer DNA methylation by statistically testing current signals between native DNA reads and methylation-free DNA reads. Tombo can detect all types of DNA methylation without priori knowledge of current signal patterns on specific methylation types. However, Tombo is not reliable at single nucleotide level and usually has high false positive rate^23^. (2) Model-based methods, such as nanopolish^24^, signalAlign^25^, DeepMod^26^, and DeepSignal^27^, utilize hidden Markov models or deep neural networks to predict specific sites as modified or unmodified. Model-based methods achieve high accuracies on 5mC detection in a specific motif, such as CpG or C*C*WGG (W indicates A or T)^28^. However, there is currently no method which can profile methylation of cytosines in both CpG and non-CpG contexts with acceptable accuracies. (3) Basecalling-based methods, such as Megalodon^29^, directly call modified bases using an extended alphabet during basecalling^23^. By using basecalling models of Guppy, which is the official ONT basecaller, Megalodon can detect a variety of methylation types, including 5mC in all contexts. However, the capability of Megalodon for non-CpG methylation detection is lack of evaluation.

Herein, we developed a pipeline for accurate and comprehensive 5mC detection in plants from native Nanopore reads by utilizing deep neural networks (DNN). First, we pair sequenced Nanopore reads and bisulfite sequencing reads of two model plants *A. thaliana* and Oryza sativa (*O. sativa*), respectively. We found that fully methylated cytosines are much less than fully unmethylated cytosines in *A. thaliana* and *O. sativa* (Results), which makes it difficult to collect positive training samples and thus results in unbalanced training dataset. Then, we developed a sample selection strategy in our pipeline to balance and denoise training samples, which significantly improved the performances of the trained models on non-CpG methylation detection. We trained three DNN models to detect 5mC sites in CpG, CHG and CHH sequence contexts, respectively. Results of the trained models in *A. thaliana* and *O. sativa* showed high agreement with bisulfite sequencing. We also tested the trained models from Nanopore reads of Brassica nigra (*B. nigra*), which also achieved high correlations with bisulfite sequencing in both CpG and non-CpG methylation detection. Furthermore, since Nanopore sequencing does not have amplification biases and has much longer read length than bisulfite sequencing, our proposed pipeline can profile more 5mC sites in plants than bisulfite sequencing, especially in highly repetitive regions.

## Results

### A novel pipeline for DNA 5mC detection in plants by Nanopore sequencing

We sequenced two model plants *A. thaliana* and *O. sativa* with bisulfite sequencing and Nanopore sequencing (Methods). By using bisulfite sequencing, we sequenced three technical replicates of *A. thaliana*: ∼116×, ∼131× and ∼116× coverage of reads, respectively. For *O. sativa*, we sequenced bisulfite sequencing data of two biological replicates: ∼78× and ∼126× coverage of reads, respectively. In *A. thaliana* and *O. sativa* genome, there are >42.8M cytosines and >162.5M cytosines, respectively. For *A. thaliana*, bisulfite sequencing detected ∼98.3% cytosines (Supplementary Table 1). We observed about 24.3% CpG, 8.6% CHG and 3.3% CHH methylation at overall genome-wide level of *A. thaliana* in all three technical replicates (Supplementary Fig. 1a). For *O. sativa*, bisulfite sequencing detected ∼93.3% and ∼94.0% cytosines in the two biological replicates (Supplementary Table 1), respectively. One replicate (rep1) has 52.7% CpG, 27.7% CHG and 4.5% CHH methylation at overall genome-wide level and the other replicate (rep2) has 46.8% CpG, 20.3% CHG and 2.9% CHH methylation at overall genome-wide level (Supplementary Fig. 2a). Compared to CpG motif, CHG and CHH motif showed lower methylation levels (Supplementary Figs. 1-2).

We further compared the number of fully unmethylated and methylated cytosines in *A. thaliana* and *O. sativa*. We observed that fully methylated cytosines are much less than fully unmethylated cytosines in both *A. thaliana* and *O. sativa*, especially in non-CpG contexts: <1:50, <1:1,000, <1:22,000 for CpG, CHG, and CHH sites in *A. thaliana*, and ∼1:2, ∼1:15 and ∼1:2,000 for CpG, CHG, and CHH sites in *O. sativa*, respectively (Supplementary Fig. 3). The huge difference of fully unmethylated and methylated cytosines in quantities, especially in non-CpG contexts, limits the collection of appropriate samples to train models for 5mC detection from Nanopore reads.

We generated ∼600× Nanopore reads of the same *A. thaliana* sample used in bisulfite sequencing (Methods). We also generated ∼215× and ∼100× Nanopore reads for the two biological *O. sativa* samples which were used in bisulfite sequencing, respectively (Methods). Then, we developed a novel pipeline which can accurately detect 5mC sites in all three contexts (CpG, CHG and CHH) from Nanopore reads (Supplementary Fig. 4). In this pipeline, we proposed DeepSignal-plant, a deep-learning method which utilizes bidirectional recurrent neural network^30^ (BRNN) with long short-term memory^31^ (LSTM) units to detect DNA 5mC methylation from Nanopore reads (Fig. 1a, Methods). For each targeted site (*i*.*e*., a sample), DeepSignal-plant requires four *k*-length features of the *k*-mer (*k*=13 by default) which the targeted site centers on it as sequence features, and *m*-length (*m*=16 by default) signals of each base in the *k*-mer to form signal features (Methods). By using BRNN to construct signal features instead of inception blocks^32^ in DeepSignal^27^, the model of DeepSignal-plant is ∼8 times smaller in size than DeepSignal (Supplementary Table 2).

**Fig 1.**
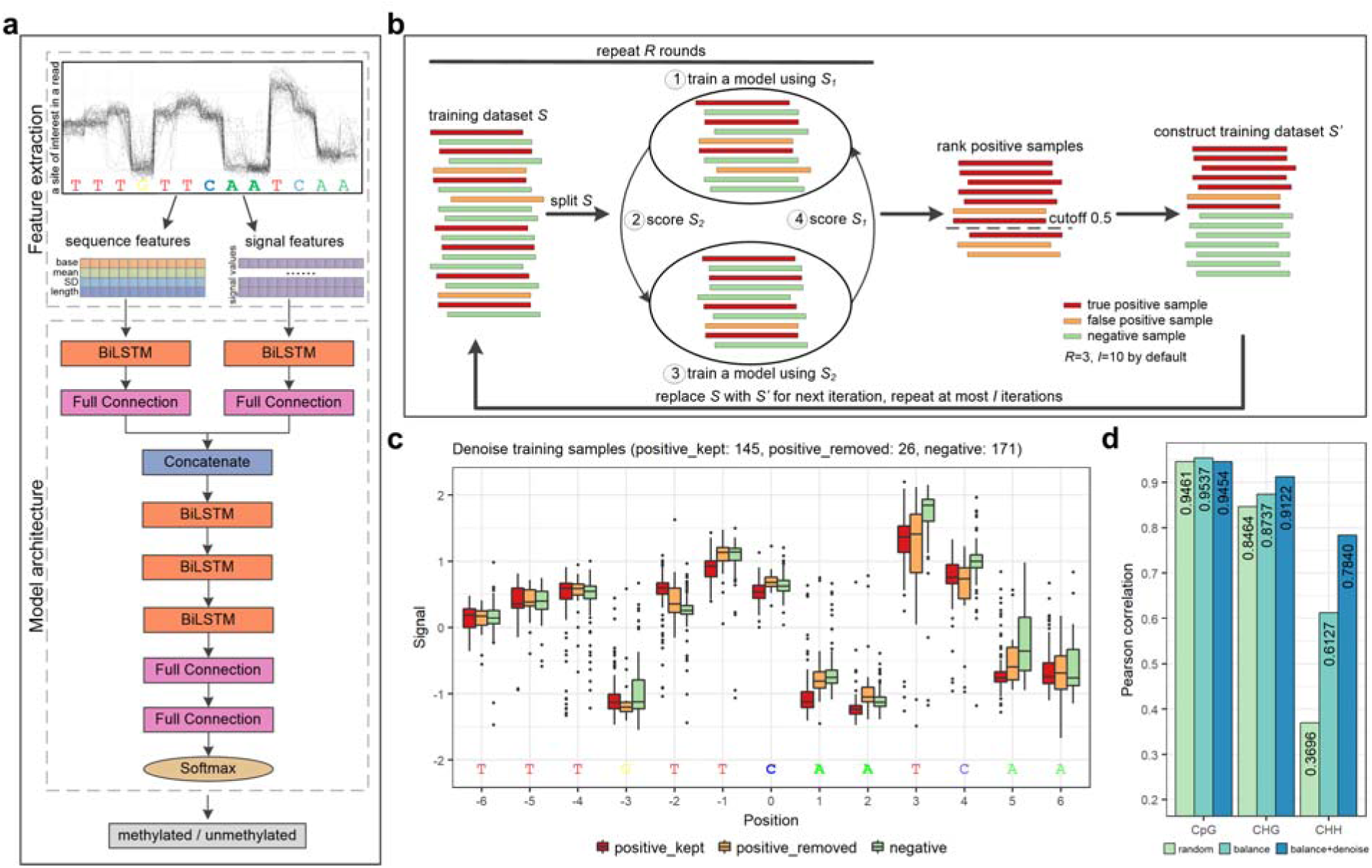
Our proposed pipeline for 5mC detection using Nanopore sequencing. **a**. Flowchart of DeepSignal-plant. BiLSTM: bidirectional long short-term memory layer; Full Connection: fully connected layer. **b:** Schematic diagram of denoise training samples using DeepSignal-plant. **c:** Signal comparison of different kinds of samples of an *k*-mer in denoising training samples. positive_kept: positive samples kept by the *denoise training samples* step; positive_removed: positive samples removed by the *denoise training samples* step; negative: negative samples. **d:** Effectiveness of training samples selection on 5mC detection. The training samples were extracted from ∼500× Nanopore reads of *A. thaliana*. Pearson correlations were calculated using the results from ∼20× Nanopore reads and three bisulfite replicates of *A. thaliana*. **e:** Cross species validation of our proposed pipeline on 5mC detection. m_arab, m_rice, m_comb represent the models of DeepSignal-plant trained using ∼500× *A. thaliana* Nanopore reads, ∼115× *O. sativa* Nanopore reads and the combined Nanopore reads, respectively. Pearson correlations are calculated using the results from ∼20× Nanopore reads of *A*.*thaliana* and *O. sativa* with the corresponding bisulfite replicates, respectively.

We used bisulfite sequencing as benchmark to train models of DeepSignal-plant: (1) *Select high-confidence sites*. The methylated and unmethylated sites with high confidence were selected based on bisulfite sequencing. We took fully unmethylated sites of which the methylation frequencies were zero and had at least five mapped reads as high-confidence unmethylated sites. To include more high-confidence methylated sites, we took sites which had at least 0.9 methylation frequency and were covered with at least five reads (Supplementary Fig. 3, Supplementary Table 3). The confidence and reliability of site selection can be further improved if multiple technical replicates of bisulfite sequencing are available (Methods). (2) *Basecall and re-squiggle*. To predict 5mC from Nanopore reads, raw reads of Nanopore sequencing need to be basecalled, and then be re-squiggled by Tombo^22^ to map raw electrical signal values to contiguous bases in genome reference (Methods). (3) *Extract, balance and denoise training samples*. The training samples of DeepSignal-plant were extracted and then randomly selected from Nanopore reads which are aligned to the selected high-confidence sites (Methods). Due to the difference of high-confidence unmethylated and methylated cytosines, there exist differences between high-confidence unmethylated and methylated *k*-mers, especially in CHG and CHH sequence contexts (Supplementary Table 4). Thus, the number of positive and negative samples of each *k*-mer need to be balanced from all extracted training samples (Methods). To include more sites and *k*-mers in positive samples, we use sites of which the methylation frequencies are at least 0.9 as high-confidence methylated sites. However, false positive samples may also be introduced. Thus, after balancing training samples, we further used DeepSignal-plant to automatically remove false positive samples (Fig. 1b, Methods), which can ensure the reliability of positive training samples, and keep the diversity of *k*-mers at the same time. After extracting and selecting training samples, three models for CpG, CHG, and CHH motif were trained separately by DeepSignal-plant (Methods).

To examine the performance of our proposed pipeline, we randomly selected ∼500× Nanopore reads of *A. thaliana* for training, and randomly selected ∼20× reads (*i*.*e*., ∼10× reads for forward and complementary strand of the genome, respectively) from the other ∼100× coverage of reads for testing (Methods). We used bisulfite sequencing as benchmark and calculated Pearson correlation between the per-site methylation frequencies predicted by bisulfite sequencing and Nanopore sequencing. We first tested the effectiveness of the training pipeline. Fig. 1c showed that, the electrical signals of the bases of the samples removed by the denoise step were closer to the signals of the negative samples, which demonstrated that the denoise step was truly capable of removing false positive samples. After the denoise step, 19.1% training samples of CHG and 29.4% training samples of CHH were removed. As shown in Fig. 1d, compared with randomly selecting samples, balancing and denoising training samples significantly improved the performance of 5mC detection, especially for CHH motif. After balancing and denoising training samples, the correlation with bisulfite sequencing increased from 0.8464 to 0.9122 for CHG motif. For CHH motif, the correlation increased from 0.3696 to 0.7840. The results demonstrated that the imbalance of positive and negative training samples for *k*-mers, along with the false positive training samples, led to the poor capability of trained models for 5mC detection. It is worth noting that the performance of CpG motif was not improved after denoising training samples. Because that for CpG motif, we took the intersection of high-confidence methylated sites from all bisulfite replicates as the final high-confidence methylated sites set. Thus, the training dataset of CpG motif is more reliable and can be used to train models without denoising.

We then performed a cross-species validation of DeepSignal-plant. For each motif, we trained models of DeepSignal-plant by using training samples from *A. thaliana* and *O. sativa* Nanopore reads independently. Then we tested the trained models on both *A. thaliana* and *O. sativa* Nanopore reads. The same as previous experiments, for *A. thaliana*, we used the selected ∼500× and ∼20× Nanopore reads for training and testing, respectively. For *O. sativa*, we used Nanopore reads of one biological replicate (sample1): ∼115× randomly selected reads for training, and ∼20× reads from the left ∼100× reads for testing (Methods). Besides using reads of one species to train, we also trained models of DeepSignal-plant for each motif by combining the training reads of *A. thaliana* and *O. sativa*. As shown in Supplementary Fig. 5, the models trained using the combined reads achieved the overall best performances. More specifically, for CpG and CHG motif, whether the models trained using reads of one species only or the models trained using the combined reads, all achieved high correlations with bisulfite sequencing on both tested data. For CHH motif, the model trained using reads of *A. thaliana* did not perform well on the tested data of *O. sativa*. It may due to the relatively less methylated sites and *k*-mers of CHH motif in *A. thaliana* (Supplementary Tables 3-4). However, the model trained using reads of *O. sativa* achieved similar performance with the model trained using the combined reads. We further randomly selected ∼20× reads of *A. thaliana* and *O. sativa* for 5 repeated times, and used the models trained using the combined reads to call methylation from the selected reads. The 5 repeated tests achieved consistent correlations with bisulfite sequencing (standard deviation<0.003), and had high correlations with each other (Supplementary Fig. 6). These results showed that the predictions of our trained models were highly reproducible. In summary, this analysis demonstrated both effectiveness and robustness of our proposed pipeline on 5mC detection in plants.

### Evaluation of the proposed pipeline

To further evaluate our proposed pipeline, we then compared DeepSignal-plant with existing methods (Tombo^22^, nanopolish^24^, DeepSignal^27^ and Megalodon^29^). For DeepSignal-plant, we used models which are trained by using the combined training samples of *A. thaliana* and *O. sativa* to call methylation from the selected reads. For 5mC detection of *A. thaliana* and *O. sativa*, DeepSignal-plant outperformed all other methods (Supplementary Fig. 7). Specifically, for 5mC detection in CpG contexts, all methods achieved high correlations with bisulfite sequencing, especially the model-based methods (nanopolish, DeepSignal, Megalodon and DeepSignal-plant). Among the compared methods, nanopolish got the closest performances with DeepSignal-plant (Supplementary Fig. 7b). It worth noting that nanopolish only gives a single prediction for coupled CpG sites on two strands of a genome by default, as nanopolish assumes that CpG sites are not hemimethylated. To compare with nanopolish, we also summed the predictions from both strands in the results of other methods. For 5mC detection in non-CpG contexts, Tombo and Megalodon, which are claimed to have the ability to detect 5mC in all contexts, were compared. However, with the built-in models or algorithm, the two methods did not get acceptable performances (Supplementary Fig. 7a).

Megalodon is a tool that provides easy-to-use interfaces to train models. We trained new models of Megalodon by using the same dataset for training models of DeepSignal-plant (Methods). Then, we used DeepSignal-plant and the newly trained models of Megalodon to detect 5mCs in ∼100× (∼50× reads for forward and complementary strand of the genome, respectively) Nanopore reads of *A. thaliana* and *O. sativa* (sample1), respectively (Methods). Then we calculated methylation frequencies of cytosines to compare with bisulfite sequencing (Methods). Compared to Megalodon, DeepSignal-plant achieved higher Pearson correlations and lower root mean square errors (RMSE) for detecting 5mCs in all contexts (Supplementary Table 5). For *A. thaliana*, DeepSignal-plant got correlation values of 0.9840 and 0.9638 with bisulfite sequencing for CpG and CHG methylation, and got a 0.8975 correlation for CHH methylation (Supplementary Fig. 8a-c). For *O. sativa*, the correlation values of DeepSignal-plant with bisulfite sequencing for CpG, CHG and CHH methylation were 0.9919, 0.9603 and 0.8563, respectively (Supplementary Fig. 8d-f). With newly trained models, Megalodon improved the performances for 5mC detection (Supplementary Fig. 9, Supplementary Table 5). For CpG methylation, Megalodon got 0.9654 and 0.9871 correlations with bisulfite sequencing in *A. thaliana* and *O. sativa*, respectively (Supplementary Fig. 9a and d). For CHG methylation, the correlations were improved to 0.9353 and 0.9551, respectively (Supplementary Fig. 9b and e). However, Megalodon did not perform well for CHH methylation, even with the newly trained models: up to 0.5783 and 0.7157 for *A. thaliana* and *O. sativa*, respectively (Supplementary Fig. 9c and f). Extra care may should be taken for Megalodon when training a model of a motif which has low methylation level. The advantage of DeepSignal-plant was greater when lower coverage of reads were used (Fig. 2a-b). With 20× coverage of reads, Megalodon got 0.9430 and 0.9776 correlations with bisulfite sequencing for CpG methylation in *A. thaliana* and *O. sativa*, whereas DeepSignal-plant achieved 0.9748 and 0.9882 correlations, respectively. For CHG methylation in two plants, DeepSignal-plant achieved ∼0.93 and ∼0.95 correlations with 20× coverage of reads, while Megalodon needed 60× coverage of reads to achieve the same correlations. For CHH methylation in *A. thaliana*, DeepSignal-plant achieved an >0.83 correlation with only 20× coverage of reads, and achieved an >0.86 correlation with 40× coverage of reads. For CHH methylation in *O. sativa*, DeepSignal-plant got a correlation value of >0.81 with 20× coverage of reads, and achieved a >0.85 correlation with 80× coverage of reads. For O. sativa, we also compared the two tools by using ∼100× Nanopore reads of the other biological replicate (sample2), which got consistent results (Supplementary Fig. 10, Supplementary Table 5). In summary, the results showed that by using our proposed pipeline, Nanopore sequencing can achieve high performance on 5mC detection in plants, even with low coverage of reads.

**Fig 2.**
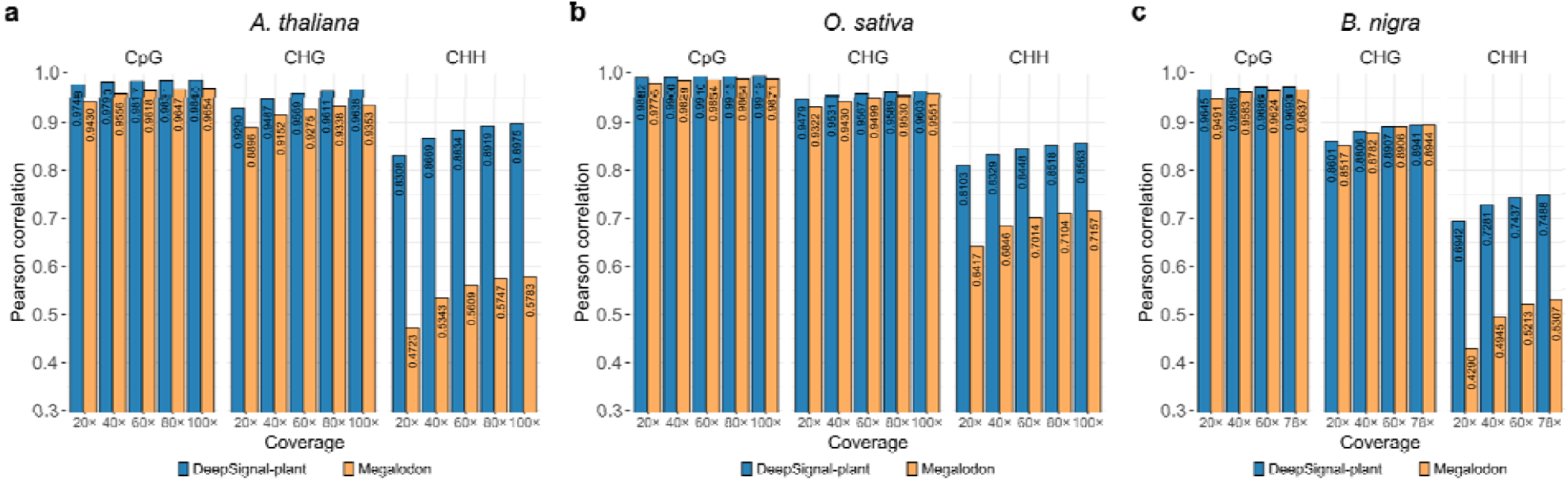
Comparison between DeepSignal-plant and Megalodon against bisulfite sequencing on 5mC detection under different coverages of Nanopore reads in *A. thaliana* (**a**), *O. sativa* (sample1) (**b**) and *B. nigra* (**c**). For each coverage (20× to 80× for *A. thaliana* and *O. sativa* (sample1), 20× to 60× for *B. nigra*), the reads were randomly shuffled and selected from the whole ∼100×/78× reads. Values for 20×, 40×, 60×, and 80× are average of 5 replicated tests.

To further assess the results of DeepSignal-plant in all three contexts, we categorized cytosines into three bins based on their methylation frequencies calculated from bisulfite sequencing: low methylation (0.0-0.3), intermediate methylation (0.3-0.7), and high methylation (0.7-1.0) (Methods). Then, we compared the spread of predictions from DeepSignal-plant and Megalodon (Supplementary Figs. 11-12). The results showed that, both DeepSignal-plant and Megalodon had a high consistency with bisulfite sequencing for predicting lowly methylated sites in all three contexts. However, Megalodon tended to underpredict highly and intermediately methylated cytosines, especially in CHH contexts, whereas the results of DeepSignal-plant were more consistent with bisulfite sequencing.

Recently Parkin *et al*. sequenced ∼78× Nanopore reads of *B. nigra* (genotype Ni100), along with ∼20× bisulfite sequencing reads^33^. A high-contiguity *B. nigra* genome (491M, 8 chromosomes) was assembled in their work. According to bisulfite sequencing, there are 70.2% CpG, 27.9% CHG and 8.3% CHH methylation at overall genome-wide level in *B. nigra* genome. We assessed DeepSignal-plant and Megalodon for 5mC detection using the Nanopore sequencing data of *B. nigra* (Fig. 2c, Supplementary Table 6). By using the whole ∼78× Nanopore reads, DeepSignal-plant got correlation values of 0.9693, 0.8941 and 0.7488 for CpG, CHG and CHH methylation, whereas the correlation values for Megalodon are 0.9637, 0.8944, and 0.5307, respectively. DeepSignal-plant and Megalodon achieved comparable correlations for CpG and CHG methylation. However, DeepSignal-plant achieved much higher correlation and lower RMSE than Megalodon for CHH methylation (Supplementary Table 6). More importantly, DeepSignal-plant were more stable under low coverage of reads (Fig. 2c). Megalodon underpredicted highly methylated sites, especially for CHH methylation, whereas DeepSignal-plant had a better performance in predicting both lowly and highly methylated sites (Supplementary Fig. 13). In summary, the results showed that models trained by our proposed pipeline can accurately detect 5mCs in cross-species Nanopore reads.

### Nanopore sequencing profiles methylation of more cytosines than bisulfite sequencing

The reads generated by Nanopore sequencing are usually 10-60 kb (kilobase) in length^34^, which are much longer than the reads of next-generation sequencing. This feature enables Nanopore sequencing to detect more cytosines that previously cannot be predicted by bisulfite sequencing. To assess this advantage of Nanopore sequencing, we then analyzed the methylation profile of cytosines detected by our proposed pipeline. In the assembled genome of *B. nigra*, there are ∼189M cytosines. By using ∼78× Nanopore reads, our proposed pipeline can profile methylation of 99.4% cytosines (Supplementary Table 7). However, we noted that the bisulfite sequencing data of *B. nigra* is only ∼20×, which is insufficient (Supplementary Table 7). Thus, we did not include *B. nigra* in this analysis.

By using Nanopore sequencing, our proposed pipeline can detect more cytosines than bisulfite sequencing with only 40× coverage of reads (Supplementary Fig. 14). With ∼100× coverage of Nanopore reads, our proposed pipeline profiled methylation of ∼99.4% and ∼99.3% cytosines (coverage>=5) in *A. thaliana* and *O. sativa* genome, respectively (Methods, Supplementary Table 1). Compared to bisulfite sequencing, our proposed pipeline detected >1.1% more cytosines in *A. thaliana* genome (Supplementary Table 1, Supplementary Fig. 15a, Supplementary Fig. 16a-c). In the two biological replicates of *O. sativa*, our proposed pipeline detected >5.9% and >5.3% more cytosines, respectively (Supplementary Table 1, Supplementary Fig. 15b-c, Supplementary Fig. 16d-i). In *A. thaliana*, among those cytosines which can only be detected by Nanopore sequencing, about 74.2% cytosines were lowly methylated and 12.8% cytosines were highly methylated (Supplementary Fig. 17a, Supplementary Fig. 18a-c). In *O. sativa*, most of the newly profiled cytosines were either lowly methylated or highly methylated: 70.4%, 24.3% cytosines were highly and lowly methylated in sample1 (Supplementary Fig. 17b, Supplementary Fig. 18d-f), and 63.5%, 29.3% cytosines were highly and lowly methylated in sample2 (Supplementary Fig. 17c, Supplementary Fig. 18g-i).

We then examined if the cytosines which can only be detected by our proposed pipeline are enriched in biological regions of *A. thaliana* and *O. sativa* genome. We found that a significant amount of cytosines were in centromeres, pericentromeres and telomeres, which are composed of thousands of repeats^35,36^ (Fig. 3a-b, Supplementary Fig. 19). In *A. thaliana*, 23.0% of these cytosines were in repeat regions (Supplementary Fig. 20a). There were 15.6%, 10.3 of these cytosines in protein-coding genes and transposons of *A. thaliana*, respectively (Supplementary Fig. 20b-c). In *O. sativa*, there were even more cytosines which can only be detected by Nanopore sequencing in repeat regions: 36.5% and 43.4% for sample1 and sample2, respectively (Supplementary Fig. 20d and g). Less newly detected cytosines were in gene regions of *O. sativa*: 6.3% and 7.3% were in protein-coding genes of sample1 and sample2, respectively (Supplementary Fig. 20). With the newly detected cytosines, our proposed pipeline profiled methylation of gaps that were previously unmappable by bisulfite sequencing (Fig. 3c-d, Supplementary Fig. 21), which made more biological regions being fully profiled (Supplementary Fig. 22). The newly profiled biological regions may provide novel insights into gene expression and transposon silencing in plants^13,37^.

**Fig 3.**
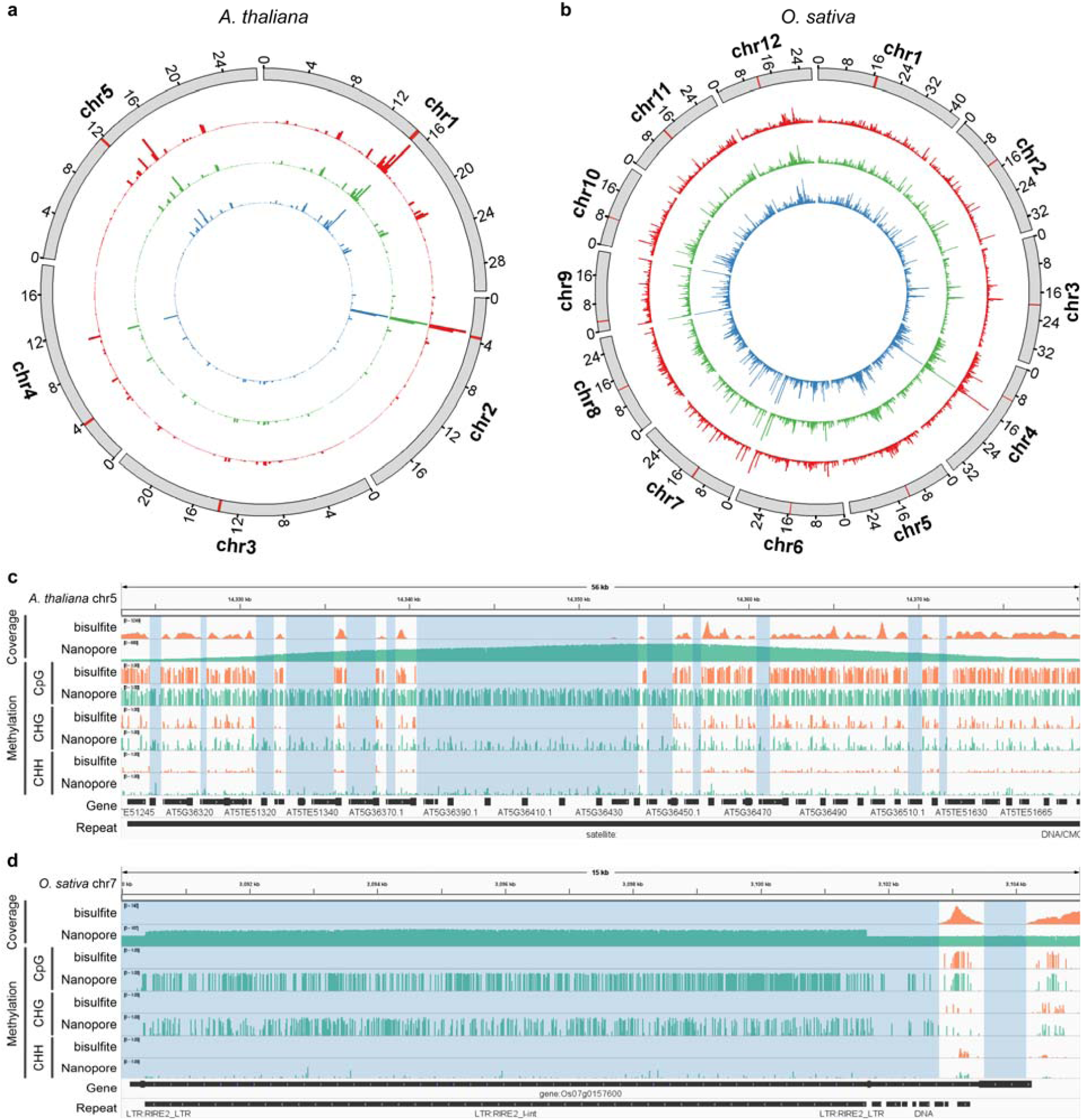
Our proposed pipeline with Nanopore sequencing profiles methylation of more cytosines than bisulfite sequencing. **a-b:** Circos plot of number of cytosines detected by Nanopore sequencing only in the genomes of *A. thaliana* (a) and *O. sativa* (sample1) (b). Cycles from inner to outer: CpG (blue), CHG (green), CHH (red), reference (the chromosomes are binned into 200,000-bp (base pair) windows. The centromeric region is indicted by red bar in each chromosome). **c-d:** Genome browser view [ref] of the reads coverage and methylation in a 56 kb region of *A. thaliana* chr5 (c) and a 14 kb region of *O. sativa* (sample1) chr7 detected by bisulfite sequencing (Bismark) and Nanopore sequencing (DeepSignal-plant). The blue shaded area shows the gaps which cannot be mapped by bisulfite sequencing.

### Methylation profiling of repeat pairs by DeepSignal-plant

We further identified repeat pairs (*i*.*e*., two same/similar sequences in genome reference) using MUMmer^38^ and used our proposed pipeline profiled to profile both CpG and non-CpG methylation in the repeat pairs (Methods). By counting differentially methylated cytosines in the repeat pairs, we found that over ∼9% and ∼6% repeat pairs in *A. thaliana* and *O. sativa*, respectively, are differentially methylated (Fig. 4a-b, Supplementary Fig. 23a, Supplementary Table 8). We also counted differentially methylated CpG and non-CpG sites in the repeat pairs (Supplementary Fig. 24, Supplementary Table 8). We found that differentially methylated repeat pairs identified by CpG, CHG, and CHH methylation independently showed largely differences^39^ (Fig. 4c-d, Supplementary Fig. 23b). CpG sites were more likely differentially methylated in repeat pairs of *A. thaliana*, while CHG sites were more likely differentially methylated in *O. sativa* (Supplementary Fig. 24). Compared to bisulfite sequencing, our proposed pipeline identified more differentially methylated repeat pairs (Supplementary Fig. 25). The differentially methylated repeat pairs in two replicates of *O. sativa* showed great inconsistency, which indicated that the differential CpG and non-CpG methylation were stable in repetitive regions (Fig. 4e, Supplementary Fig. 26). Moreover, we also found several long repeat pairs which were differentially methylated (Fig. 4f-g, Supplementary Fig. 27, Supplementary Table 8), which indicated that both CpG and non-CpG methylation may be further used to improve assembly of long repetitive regions by Nanopore sequencing.

**Fig 4.**
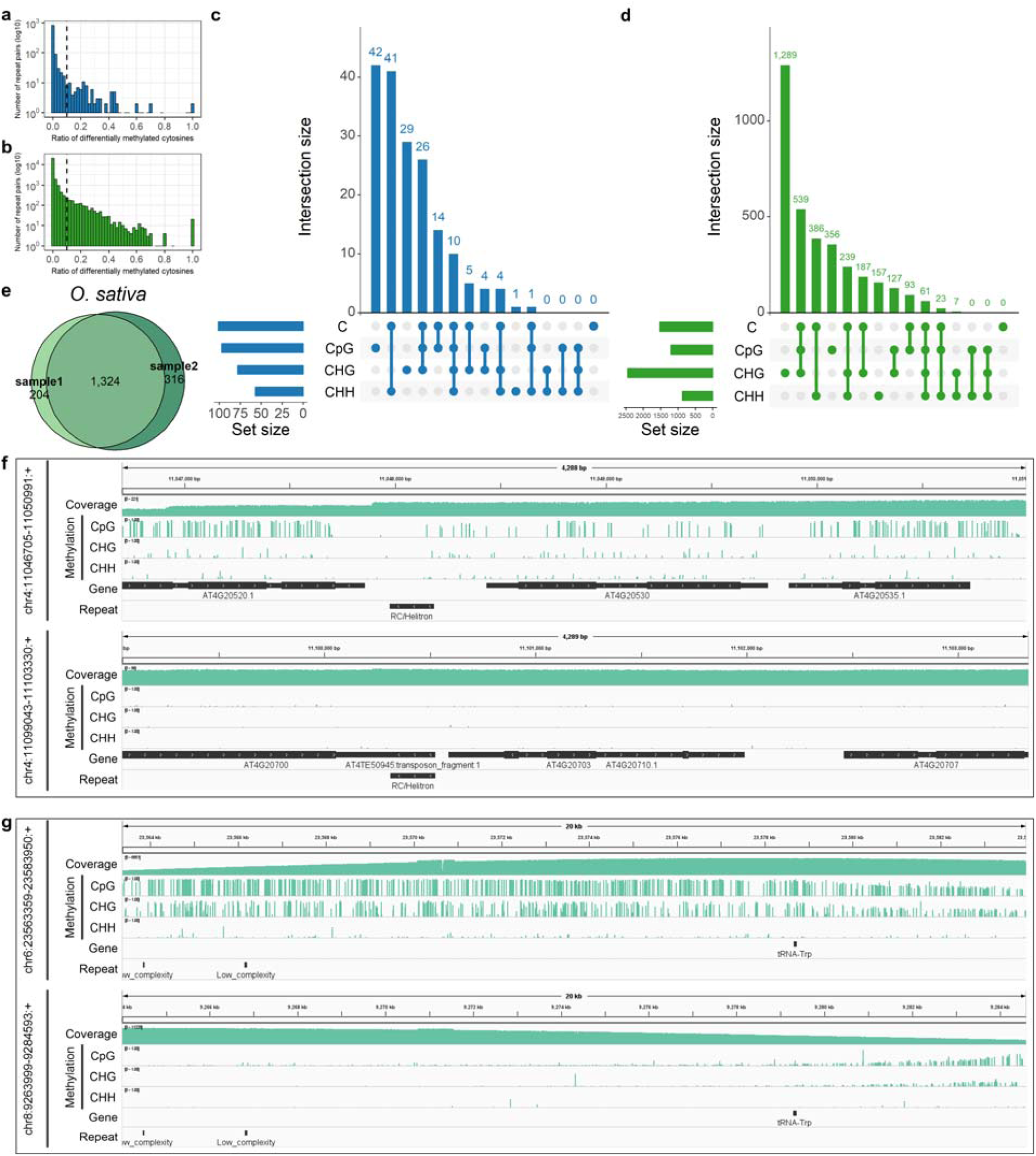
Our proposed pipeline identified differentially methylated repeat pairs. **a-b:** Ratio of differentially methylated cytosines in repeat pairs of *A. thaliana* (a) and *O. sativa* (sample1) (b). **c-d:** Matrix layout for all intersections of four sets of differentially methylated repeat pairs profiled by cytosines, CpG sites, CHG sites, and CHH sites independently, in *A. thaliana* (c) and *O. sativa* (sample1) (d). Circles in the matrix indicate sets that are part of the intersection. **e:** Comparison of differentially methylated repeat pairs identified by methylation of cytosines in *O. sativa* sample1 and sample2. **f:** Genome browser view of a differentially methylated repeat pair (chr4:11046705-11050991:+, chr4: 11099043-11103330:+) in *A. thaliana*. **g:** Genome browser view of a differentially methylated repeat pair (chr6:23563359-23583950: +, chr8: 9263999-9284593:+) in *O. sativa* (sample1).

## Discussion

In this study, we proposed a novel pipeline to detect DNA 5mC in plants from native Nanopore reads. By selecting appropriate training samples, we trained models which can accurately detect 5mCs not only in CpG sites, but also in CHG and CHH sites. Experiments on two model plants (*A. thaliana* and *O. sativa*) and *B. nigra* showed that, our proposed pipeline had high agreement with bisulfite sequencing for the predictions of CpG, CHG and CHH methylation. Our proposed pipeline can detect 5mC sites with acceptable accuracies even with low coverage of reads. Furthermore, we demonstrate that by using Nanopore sequencing, our proposed pipeline can profile methylation of more cytosines in plants than bisulfite sequencing, especially in highly repetitive genome regions. With longer reads, Nanopore sequencing is promising to have huge advantages in repetitive plant and polyploidy plant genomes. To summarize, our proposed pipeline may become a new method for 5mC detection, which will provide novel and deeper insights into the epigenetic mechanisms of plants.

Currently, only models for 5mC, the most common and well-studied DNA methylation, are provided by DeepSignal-plant in our proposed pipeline. With new ground-truth data of different modifications generated, we expect that accurate models to predict other modifications could be trained. Besides DNA modifications, Nanopore reads of direct RNA sequencing also contains signals of RNA modifications. However, to detect RNA modifications using supervised machine learning, accuracy of basecalling tools for RNA sequencing reads needs to be improved. Moreover, large-scale gold standard data of RNA modifications needs to be generated. Overall, with more training data of other modifications available in the future, we expect that DeepSignal-plant, together with our proposed pipeline, can become a powerful tool in epigenetics and epitranscriptomics analysis.

## Methods

### Plant materials and DNA extraction

Wild type Arabidopsis thaliana (L.) Heynh. Columbia-0 (Col-0) was used in this study. Seeds were surface-sterilized in 4% sodium hypochlorite, vernalized for 2 days at 4°C and grown on half Murashige & Skoog (MS) plates for 7days at 22□, 70% relative humidity with a 16 h/8 h light/dark regime. 7-day-old seedlings were then transplanted into individual pots with soil and cultivated for one month in the same condition as seedlings on half MS plates. For *O. sativa*, we used wild type Oryza sativa L. ssp. Japonica cv. Nipponbare. Seeds were soaked in distilled water for 48 hours at 37□ to accelerate germination. Then the germinated seeds were sown in soil and cultured for one month in a growth chamber at 28□, 70% relative humidity with a 14 h/10 h light/dark regime. As for taking samples of *O. sativa*, each seedling was divided into two halves in the vertical direction with scissors, marked with “left group” and “right group”. For *A. thaliana* and *O. sativa*, the harvested material was from at least 30 individual plants and quickly froze with liquid nitrogen, stored at −80□ until further use. Genomic DNA was extracted from samples by QIAGEN® Genomic DNA extraction kit (Cat#13323) according to the manufacturer’s standard operating procedure. The extracted DNA was detected by NanoDrop^™^ One UV-Vis spectrophotometer (Thermo Fisher Scientific) for DNA purity. Then Qubit® 3.0 Fluorometer (Invitrogen) was used to quantify DNA accurately.

### Bisulfite sequencing

The extracted genomic DNA was first sheared by Covaris and purified to 200-350 bp in average size. Sheared DNA was then end-repaired and ligated to methylated sequencing adapters. Finally, adapter-ligated DNA was treated with bisulfite and PCR-amplified. The libraries of three technical replicates of *A. thaliana* were sequenced on a NovaSeq6000 sequencer (Illumina) to obtain pair-end 150 bp (base pair) reads. ∼116×, ∼131× and ∼116× coverage of reads for each replicate were generated. For *O. sativa*, libraries of two biological replicates were sequenced: one biological replicate (rep1) was sequenced on a MGI2000 (BGI) sequencer to obtain pair-end 100 bp reads (∼78× coverage of reads); the other replicate (rep2) was sequenced on a NovaSeq6000 sequencer (∼126×). The sequencing reads were then processed by the standard pipeline of Bismark^40^ (v0.20.0). For each detected cytosine in CpG, CHG and CHH motif, Bismark outputs a methylation call for each of its mapped reads. Then the methylation frequency of the cytosine is calculated, which is the number of mapped reads predicted as methylated divided by the number of total mapped reads.

### Nanopore sequencing

The extracted genomic DNA was qualified, size-selected using the BluePippin system (Sage Science). Then, the genomic DNA was end-repaired and PCR adapters supplied in ONT sequencing kit (SQK-LSK109) were ligated to the end-repaired DNA. Finally, Qubit® 3.0 Fluorometer (Invitrogen) was used to quantify the size of library fragments. To generate Nanopore reads, the prepared libraries are loaded into flow cells (R9.4, FLO-PRO002) of a PromethION sequencer (ONT). Raw Nanopore reads were then basecalled by Guppy (version 3.6.1+249406c), the official basecaller of Oxford Nanopore Technologies, with *dna_r9*.*4*.*1_450bps_hac_prom*.*cfg*. In total, there were 3,124,608 reads with an average length of 23,751 bp (∼600×) for *A. thaliana*. For *O. sativa*, reads for each of the two biological replicates of were generated: 3,274,036 reads with an average length of 25,990 bp (∼215×) for rep1 and 1,671,237 reads with an average length of 23,790 bp (∼100×) for rep2, respectively.

For *A. thaliana*, we randomly selected ∼500× Nanopore reads for training. The left ∼100× reads were used for evaluation. For *O. sativa*, ∼115× reads of one replicate (rep1) were randomly selected for training. The left ∼100× reads, together with all ∼100× Nanopore reads of the other replicate (rep2) were used for evaluation.

### Genome references and annotations

The genome reference of *A. thaliana* was downloaded from NCBI with the version GCF_000001735.4_TAIR10.1^41^. The gene annotation and centromere location of *A. thaliana* were downloaded from Araport11^42^. The genome reference and gene annotation of *O. sativa* were downloaded from EnsemblPlants with the version IRGSP-1.0^43^ (Assembly GCA_001433935.1). Locations of centromeres in *O. sativa* were downloaded from Rice Annotation Project Database^44^. Repeat regions of *A. thaliana* and *O. sativa* were downloaded from NCBI Genome Data Viewer with the “RepeatMasker” track in corresponding genomes^41,45^. The tandem repeats and inverted repeats were generated by Tandem Repeats Finder^46^ (version 4.09) and Inverted Repeats Finder^47^ (version 3.05) with corresponding genome references and suggested parameters, respectively.

### Identify repeat pairs

To identify repeat pairs (*i*.*e*., two same/similar sequences in genome reference), we first used MUMmer^38^ (version 4.0.0beta2) to align the genome reference to itself. Then, from the results of MUMmer, we used custom Python scripts to select two regions of which length>=100 and identity score>=0.99 as repeat pairs. Finally, we kept repeat pairs which contains at least one cytosine for analysis. Suppose the methylation frequencies of a cytosine in the same relative position of the repeat pair are *rmet*_*1*_ and *rmet*_*2*_, the cytosine is said to be differentially methylated if |*rmet*_*1*_-*rmet*_*2*_|>=0.5. A repeat pair is said to be differentially methylated if the percent of differentially methylated cytosines (or CpG sites, CHG sites, CHH sites independently) is greater than 0.1.

### Select high-confidence sites from bisulfite sequencing

We take bisulfite sequencing as the gold standard to train models for 5mC detection from Nanopore reads. From the results of bisulfite sequencing, we took sites which were covered with at least five reads and had at least 0.9 methylation frequency as high-confidence methylated sites. Sites which had at least five mapped reads and zero methylation frequency were selected as high-confidence unmethylated sites. For CpG motif of *A. thaliana*, we took the intersection of high-confidence sites from all 3 technical replicates as the final high-confidence sites set to train models. For CHG and CHH motif, we took the sites of which the methylation frequencies are zero in all bisulfite replicates as the final high-confidence unmethylated sites. The methylation level of CHG and CHH sites are relatively lower in *A. thaliana*. Thus, we took the union of the high confidence methylated sites from 3 replicates as the final high-confidence methylated sites of CHG and CHH motif to train models (Supplementary Table 3). For *O. sativa*, high-confidence sites from rep1 were selected to train models (Supplementary Table 3).

### Framework of DeepSignal-plant

DeepSignal-plant is a deep-learning tool which utilizes BRNN with LSTM units. To infer methylation states of DNA bases, raw reads of Nanopore sequencing and a reference genome are needed. Before using DeepSignal-plant to call methylation, raw reads must be pre-processed (Fig. 1a) as follows:

1. *Basecall*. We use Guppy (version 3.6.1+249406c) for the basecalling of all the Nanopore reads.
2. *Re-squiggle*. We use Tombo^22^ (version 1.5.1) to map raw signals of reads to contiguous bases in the genome reference. In Tombo, minimap2^48^ (version 2.17-r941) is used for the alignment between reads and genome reference. Tombo corrects the insertion and deletion errors in Nanopore reads and re-annotates raw signals to match the genomic bases.

After pre-processing of raw reads, DeepSignal-plant can be used to train models and call methylation (Fig. 1a) as follows:

1. *Extract features*. After re-squiggle, raw signals of each read are normalized by using median shift and median absolute deviation (MAD) scale first^27^. Then, for each base in one read, we can get the set of normalized signals mapped to the base. Therefore, for each targeted site, we can use the *k*-mer where the targeted site centers on it and the corresponding normalized signals to form two groups of feature vectors: (1) *Sequence features*. For each base in the *k*-mer, we calculate the mean, standard deviation, and the number of its mapped signal values. Thus, we construct a *k*×4 matrix as sequence features, where there are 4 features for each base of the *k*-mer: the nucleotide base, the mean, standard deviation, and the number of signal values of each base, respectively. (2) *Signal features*. We sample *m* signals from all signals of each base to form a *k*×*m* matrix as signal features. For each base, if the number of signals is less than *m*, we paddle with zeros. We set *k*=13 and *m*=16 as default to extract features of each targeted site (Supplementary Note 2).
2. *Model architecture*. BRNN^30^ with LSTM units^31^ is used in the model of DeepSignal-plant (Fig. 1b). In detail, sequence features and signal features, are each fed into a BRNN layer (Supplementary Note 1), followed by a fully connected layer. Then, the concatenated features are received by BRNN with three hidden layers. Finally, after formation of two fully connected layers, a softmax activation function (Supplementary Note 1) is used to output two probabilities *P*_*m*_ and *P*_*um*_ (*P*_*m*_+*P*_*um*_=1), which represent the probabilities of methylated and unmethylated, respectively. The model architecture of DeepSignal-plant is optimized from the model architecture of DeepSignal^27^, in which we use BRNN instead of inception blocks^32^ to process signal features. Compared to DeepSignal, the model architecture of DeepSignal-plant has ∼8 times less parameters (Supplementary Table 2).
3. *Train models*. To train a model of DeepSignal-plant, the selected training samples were split into two datasets for training and validation. We use Adam optimizer^49^ to learn model parameters on the training dataset by minimizing the loss calculated by cross-entropy (Supplementary Note 2). The model parameters which get the best performance on the validation dataset are saved. To prevent overfitting, we use two strategies. First, we use dropout layers^50^ in LSTM layers and fully connected layers. Second, we use early stopping^51^ during training. The model parameters with current best performance on validation dataset are saved in every epoch. If the best performance of the current epoch decreases, we stop the training process. Hyperparameter tuning of DeepSignal-plant was also performed (Supplementary Note 2). According to the results, the *k*-mer length in DeepSignal-plant was set to 13 (Supplementary Fig. 28). The number of signals used to construct signal features was set to 16 (Supplementary Fig. 29). And models using both sequence and signal features to call methylation got the best performances (Supplementary Fig. 30b).
4. *Call methylation and calculate methylation frequency*.. For a targeted site in a read, DeepSignal-plant outputs the methylated probability *P*_*m*_ and unmethylated probability *P*_*um*_. If *P*_*m*_ > *P*_*um*_, the site is called as methylated, otherwise is called as unmethylated. Then, by counting the number of reads where the site called as methylated and the total reads mapped to the site, DeepSignal-plant calculates a methylation frequency of the site.

DeepSignal-plant was implemented in Python3 and PyTorch (version 1.2.0).

### Balance training samples of each *k*-mer

To train a model of DeepSignal-plant, we first extract training samples from all reads aligned to the high-confidence sites. Then, as in DeepSignal^27^, we randomly subsample at most 20 million (half positive and half negative) samples as the training dataset. However, there may exist big differences between the *k*-mers in the selected high-confidence methylated and unmethylated sites (Supplementary Table 4). This kind of difference leads to unbalanced positive and negative samples for each *k*-mer in the training dataset, which further influences model training, especially for CHH motif. Thus, to train a model with higher performance, we further balance the number of positive and negative samples of each *k*-mer. Specifically, for each *k*-mer in the training dataset, we subsample at most the same number of negative samples as positive samples. We further randomly select a certain amount of negative samples of the *k*-mers that do not have positive samples, to balance the total number of positive and negative samples. The algorithm is as follows:

**Algorithm 1:**
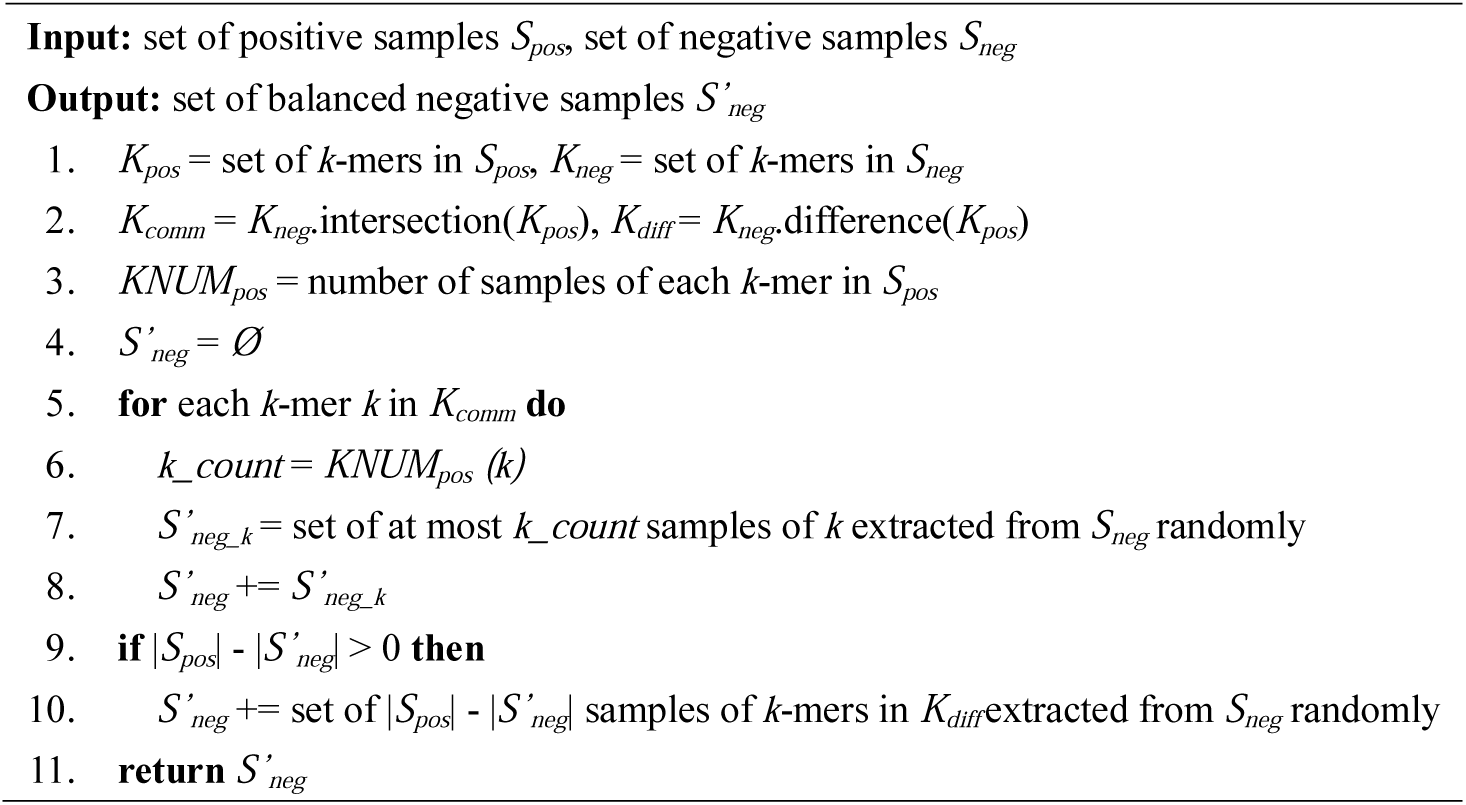
Balance_Negative_Samples (*S*_*pos*_, *S*_*neg*_)

### Denoise training samples

There exist false positive samples in the training dataset which are extracted from the reads aligned to high-confidence sites. We then design an algorithm which can iteratively remove false positive samples in the training dataset using DeepSignal-plant (Fig. 1b). In each iteration of the algorithm, we perform two-fold cross predication of the training dataset. First, the training dataset is randomly divided into two equal sets. Then the two equal sets are train two models of DeepSignal-plant separately. And the two trained models are used to predict samples of each other. For each sample, DeepSignal-plant outputs a methylation probability which ranges from 0 to 1. Thus, after cross prediction, we can rank the positive samples based on the probabilities calculated by DeepSignal-plant. To reduce the bias of cross prediction, we repeat the two-fold cross predication multiple times. At last, we remove positive samples which are predicted as unmethylated by DeepSignal-plant. The kept positive samples, together with selected negative samples are then used for next iteration. The algorithm will stop either after 10 iterations, or if there are not enough positive samples predicted unmethylated, which is described as follows:

**Algorithm 2:**
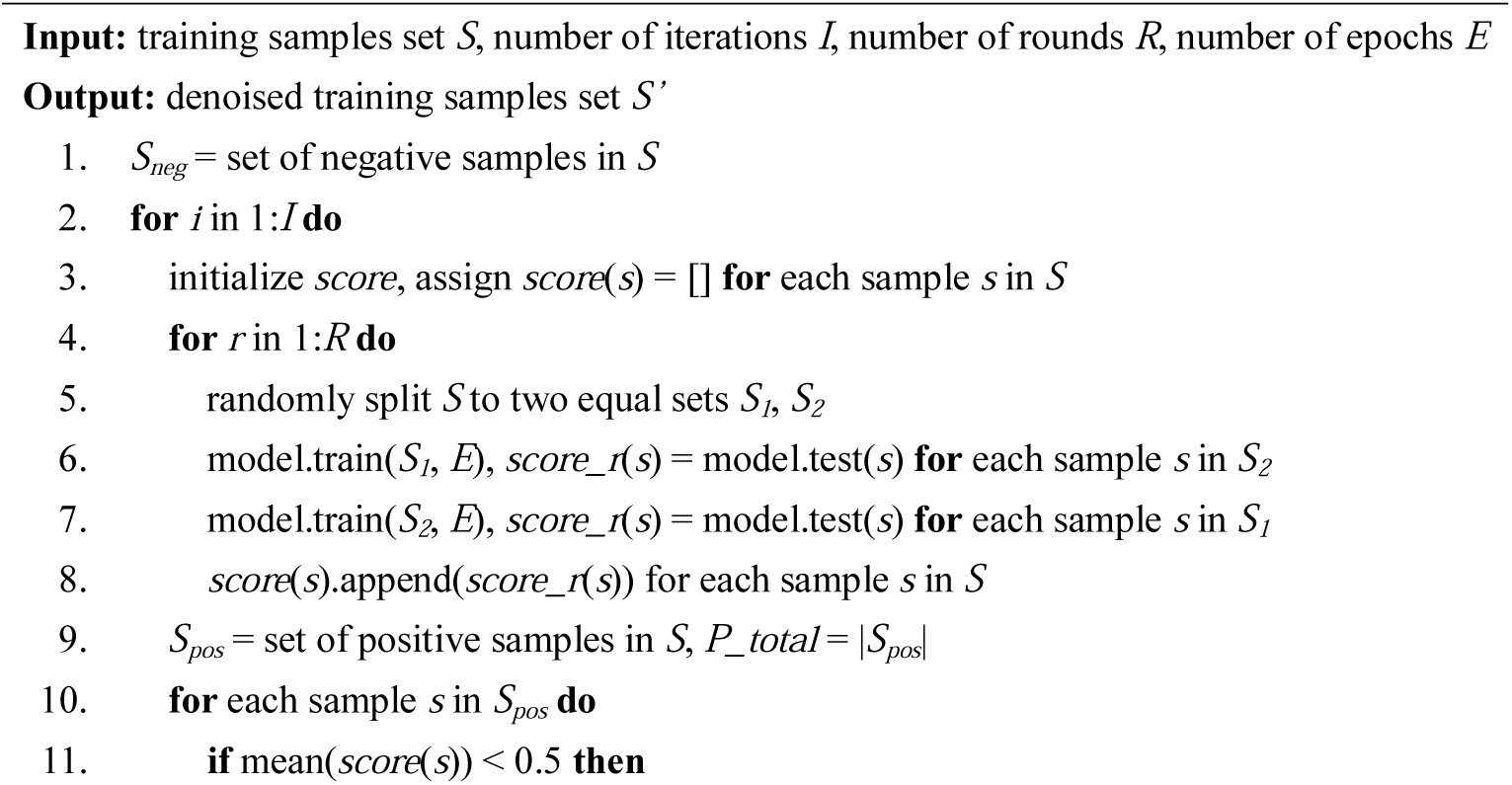
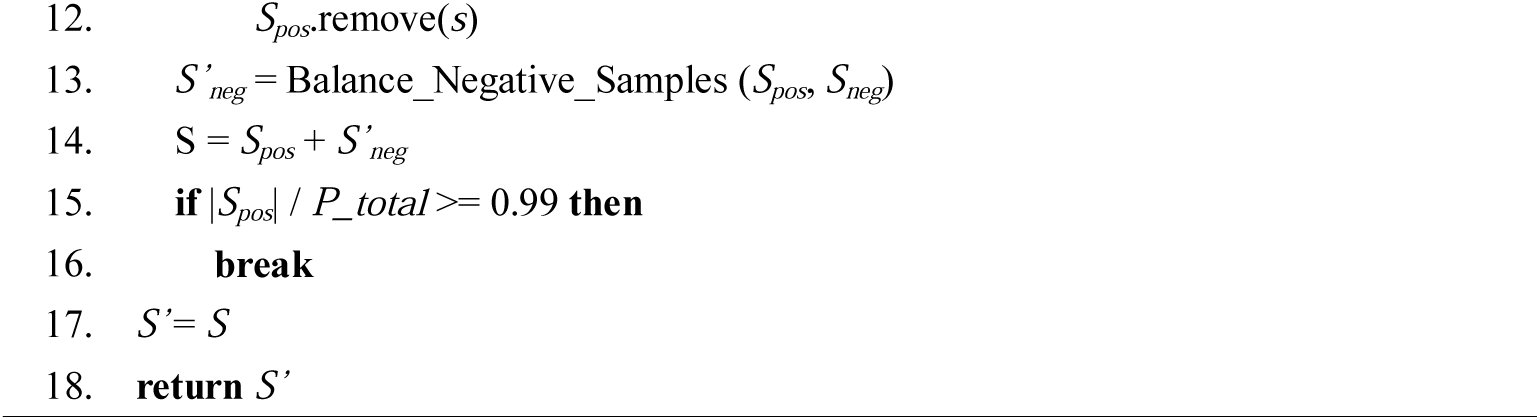
Denoise_Samples (*S, I, R, E*)

In this algorithm, we set *I*=10, *R*=3, *E*=3 as default. We used this algorithm to denoise training samples of CHG and CHH motif. According to our experiments, using only signal features to denoise training samples got the best performance (Supplementary Fig. 30a).

### Evaluation of the proposed pipeline on *A. thaliana* and *O. sativa*

We use bisulfite sequencing as benchmark to test the trained models of DeepSignal-plant. To compare with bisulfite sequencing, we used cytosines from both forward and complementary strand of genomes of *A. thaliana* and *O. sativa*. 5 chromosomes of *A. thaliana* and 12 chromosomes of *O. sativa* are used. (1) *Comparison of number of cytosines detected by bisulfite and Nanopore sequencing*. For comparison, we count sites which have at least 5 mapped reads in Nanopore sequencing and bisulfite sequencing, respectively. In bisulfite sequencing of *A. thaliana*, we count sites which have at least 5 mapped reads in at least one technical replicate. (2) *Comparison of methylation frequencies*. We use Pearson correlation (*r*), together with coefficient of determination (*r*^*2*^), Spearman correlation (ρ) and root mean square error (RMSE), to compare per-site methylation frequencies calculated by Nanopore sequencing and the corresponding replicates of bisulfite sequencing. Cytosines with at least 5 mapped reads in both bisulfite and Nanopore sequencing are selected for evaluation. For *A. thaliana*, we calculate the average correlations between Nanopore sequencing and three bisulfite replicates. To calculate methylation frequencies with different coverage of Nanopore reads, we randomly select reads from all testing reads of *A. thaliana* and *O. sativa*. (3) *Comparison of lowly, intermediately, and highly methylated sites*. A site is said to be lowly methylated if it has at least 5 mapped reads and the methylation frequency of the site is at most 0.3. A site is highly methylated if it has at least 5 mapped reads and the methylation frequency of the site is at least 0.7. The cytosines with methylation frequencies between 0.3 and 0.7 and at least 5 mapped reads are categorized as intermediately methylated sites. For Nanopore sequencing, we categorized the cytosines based on the methylation frequencies predicted from the ∼100× reads selected. For bisulfite sequencing of *A. thaliana*, we count a site as lowly, intermediately, or highly methylated if the site is lowly, intermediately, or highly methylated in all three replicates.

### Evaluation of the proposed pipeline on *B. nigra*

We got ∼78× Nanopore reads and the de novo assembly (Bnigra_NI100.v2.genome.fasta) of *B. nigra* Ni100 from Parkin *et al*.^33^. The cytosine methylation profile from bisulfite sequencing was also provided by Parkin *et al*., which was generated by BSMAP^52^ (v.2.9) from ∼20× bisulfite sequencing reads. Cytosines from both forward and complementary strand of 8 chromosomes of *B. nigra* were included for evaluation. To compare methylation frequencies with bisulfite sequencing, cytosines which have at least 5 mapped reads in Nanopore sequencing were selected. Since the coverage of bisulfite sequencing is insufficient, cytosines with at least 10 mapped reads in bisulfite sequencing were further selected.

### Comparison with existing methods

We used Tombo^22^ (version 1.5.1), nanopolish^24^ (v0.13.2), DeepSignal^27^ (v0.1.8) and Megalodon^29^ (version 2.2.3) with respective built-in models (or algorithm) for comparison. In Megalodon, two configuration files *res_dna_r941_prom_modbases_5mC_CpG_v001*.*cfg* and *res_dna_r941_min_modbases-all-context_v001*.*cfg* were used to detect 5mCs in CpG and non-CpG contexts, respectively. With the built-in models as initial models, we got new models of Megalodon for each motif (CpG, CHG and CHH) by training 2 rounds with the selected Nanopore reads of *A. thaliana* and *O. sativa*, respectively.

## Data availability

All sequencing data generated in this study (bisulfite sequencing and Nanopore sequencing data of *A. thaliana* and *O. sativa*) have been deposited in the Genome Sequence Archive of BIG Data Center, Beijing Institute of Genomics (BIG, http://gsa.big.ac.cn), Chinese Academy of Sciences, with Project Accession No. “PRJCA004326”.

## Code availability

DeepSignal-plant and a detailed tutorial are publicly available at https://github.com/PengNi/deepsignal-plant. Code for reproducing results and analysis was documented at https://github.com/PengNi/plant_5mC_analysis.

## Acknowledgements

We thank Prof. Parkin and her colleagues for sharing their bisulfite sequencing and Nanopore sequencing data of *B. nigra*. This work was supported in part by the National Natural Science Foundation of China under Grants (Nos. 61772557, 61732009 and 61828205), 111 Project (No. B18059), Hunan Provincial Science and Technology Program (No. 2018wk4001) to J.W., the U.S. National Institute of Food and Agriculture (NIFA) under grant 2017-70016-26051 and the U.S. National Science Foundation (NSF) under grants ABI-1759856 to F.L.

## Author contributions

## Supplementary information

