## Supplementary Material for "Genome-wide Detection of Cytosine Methylations in Plant from Nanopore sequencing data using Deep Learning"

### Supplementary Information

#### Supplementary Figures

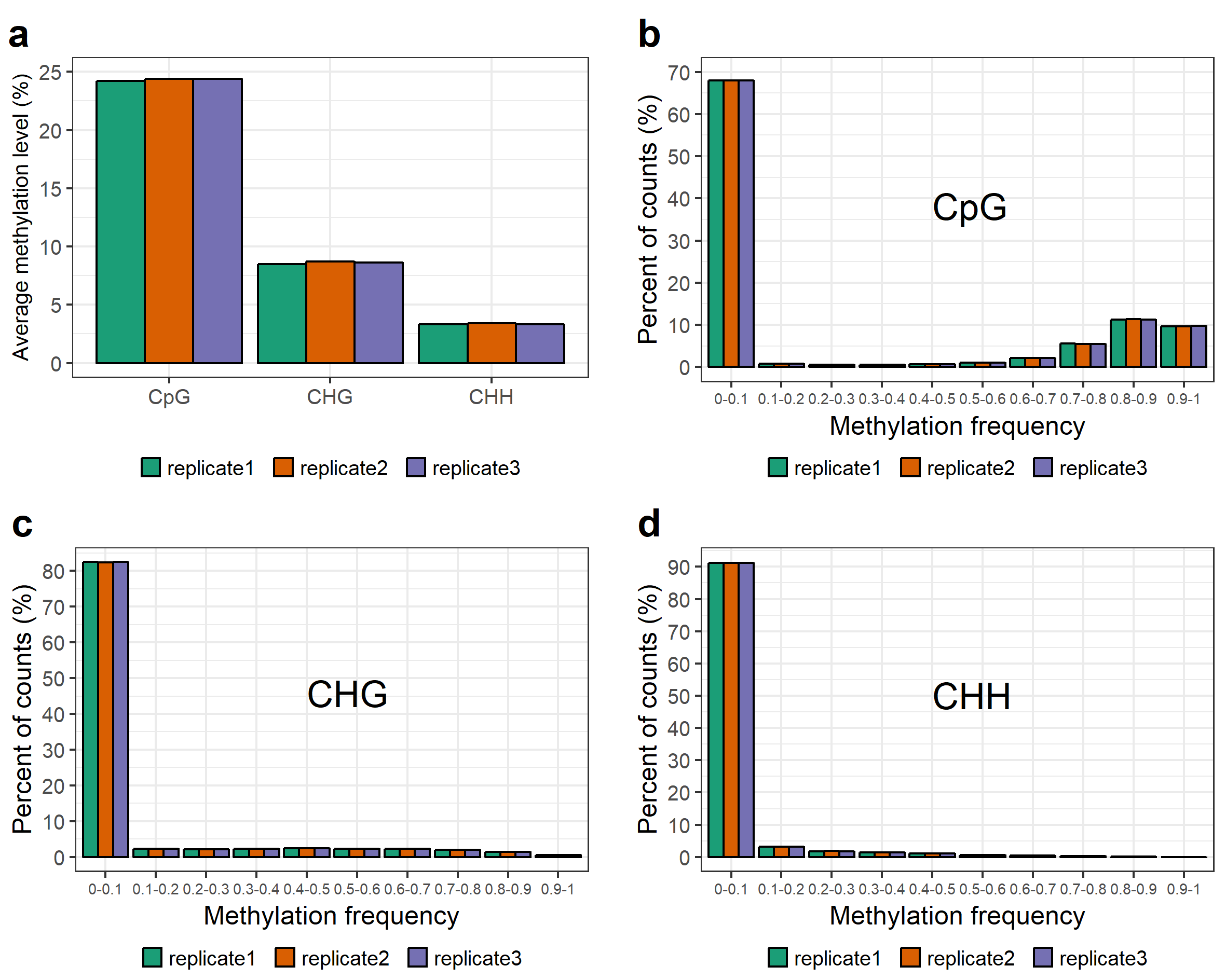

**Supplementary Fig. 1** General pattern of 5mC methylation in *A. thaliana* from bisulfite sequencing (3 technical replicates: replicate1, replicate2, and replicate3). **a:** Genome average levels of 5mC (CpG, CHG, CHH) methylation in *A. thaliana*. **b-d**: Distribution of methylation frequency of CpG (b), CHG (c) and CHH (d). The x-axis is divided into 10 bins. The y-axis is the percent of total counts for each bin respectively. When calculating methylation frequency, only the sites with at least 5 mapped reads are considered.

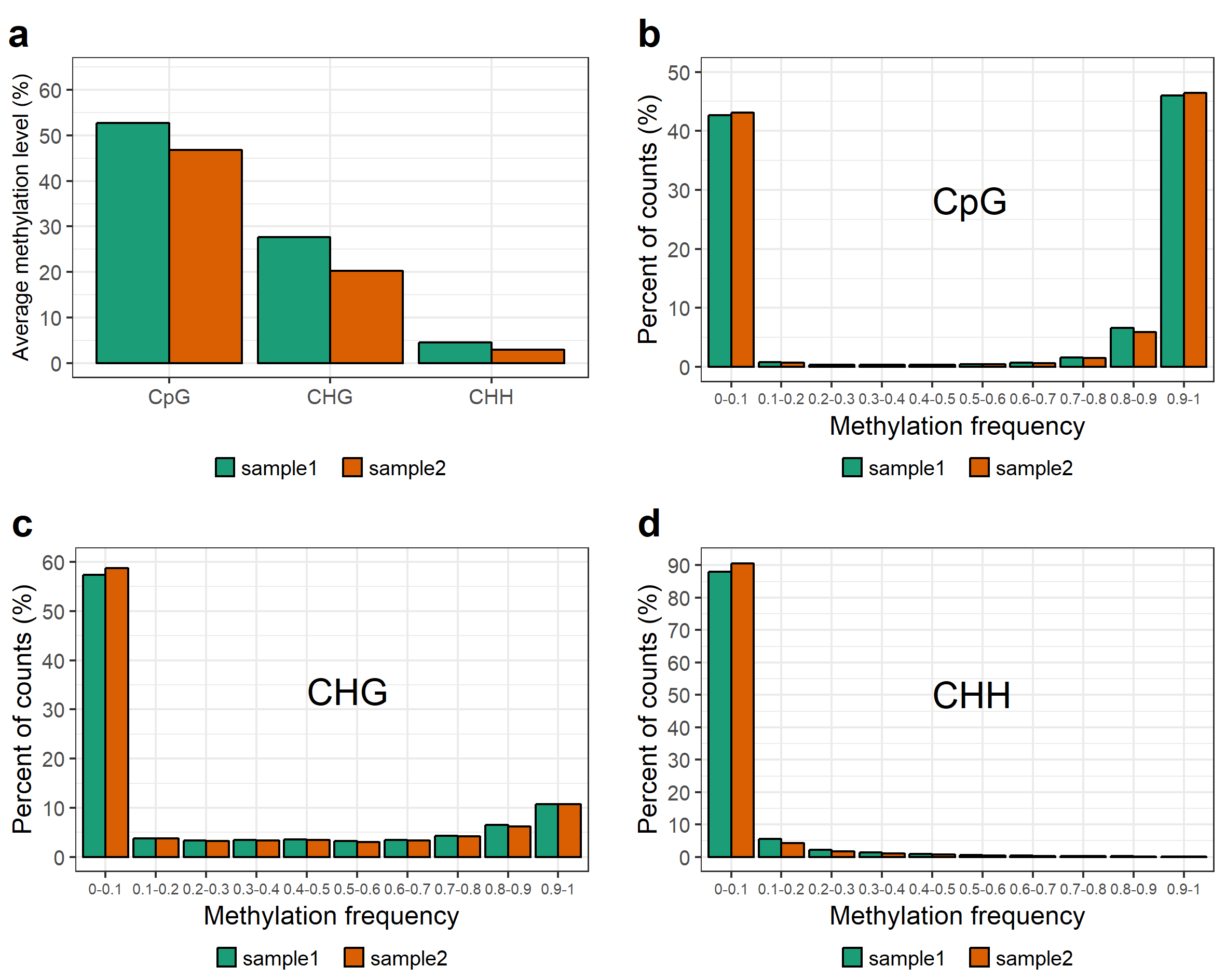

**Supplementary Fig. 2** General pattern of 5mC methylation in *O. sativa* from bisulfite sequencing (2 biological replicates: sample1 and sample2). **a:** Genome average levels of 5mC (CpG, CHG, CHH) methylation in *O. Sativa*. **b-d**: Distribution of methylation frequency of CpG (b), CHG (c) and CHH (d). The x-axis is divided into 10 bins. The y-axis is the percent of total counts for each bin respectively. When calculating methylation frequency, only the sites with at least 5 mapped reads are considered.

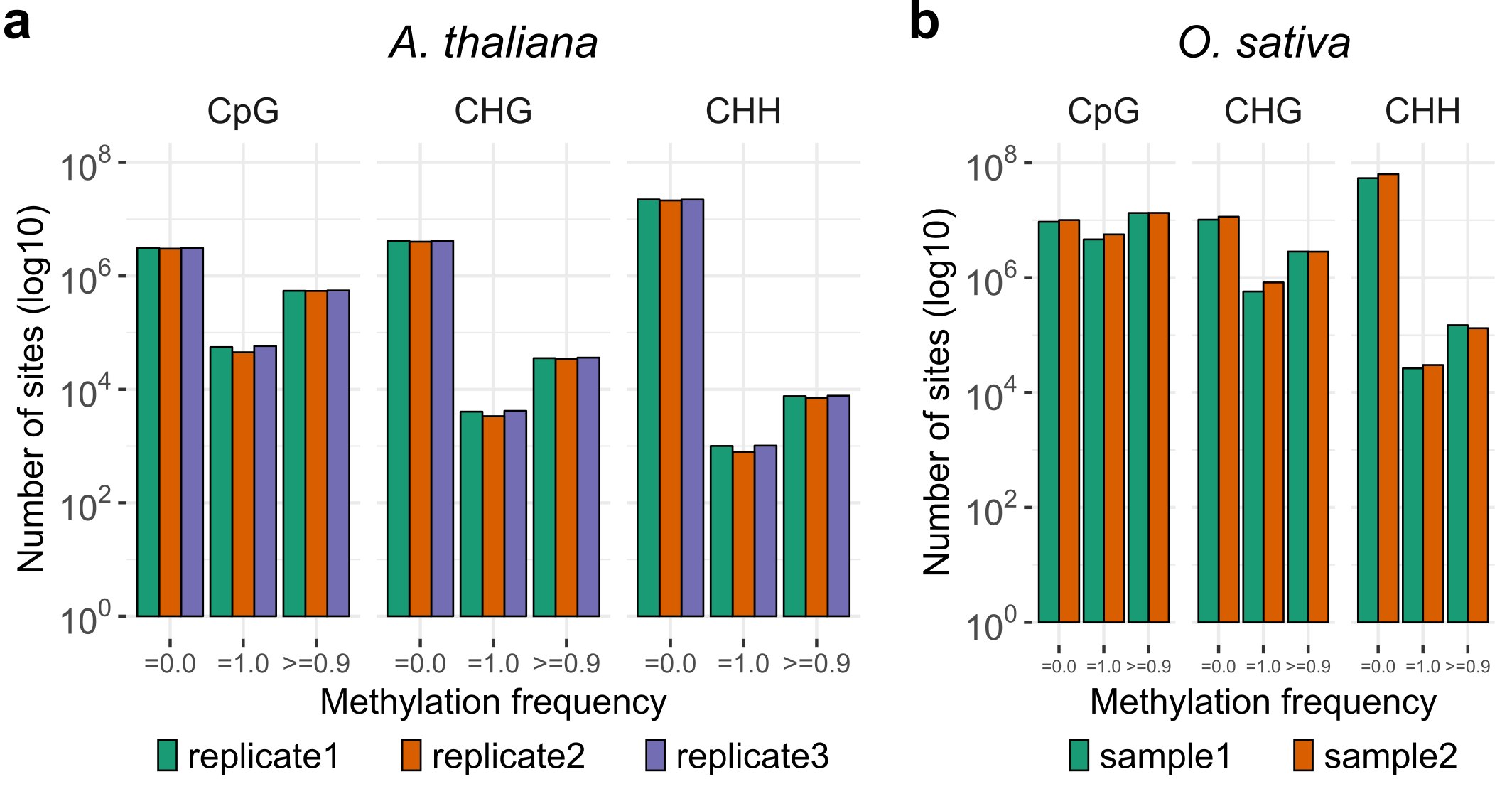

**Supplementary Fig. 3** Number of fully unmethylated sites, fully methylated sites, sites of which methylation frequency>=0.9 based on bisulfite sequencing. **a:** *A. thaliana*. **b:** *O. sativa*.

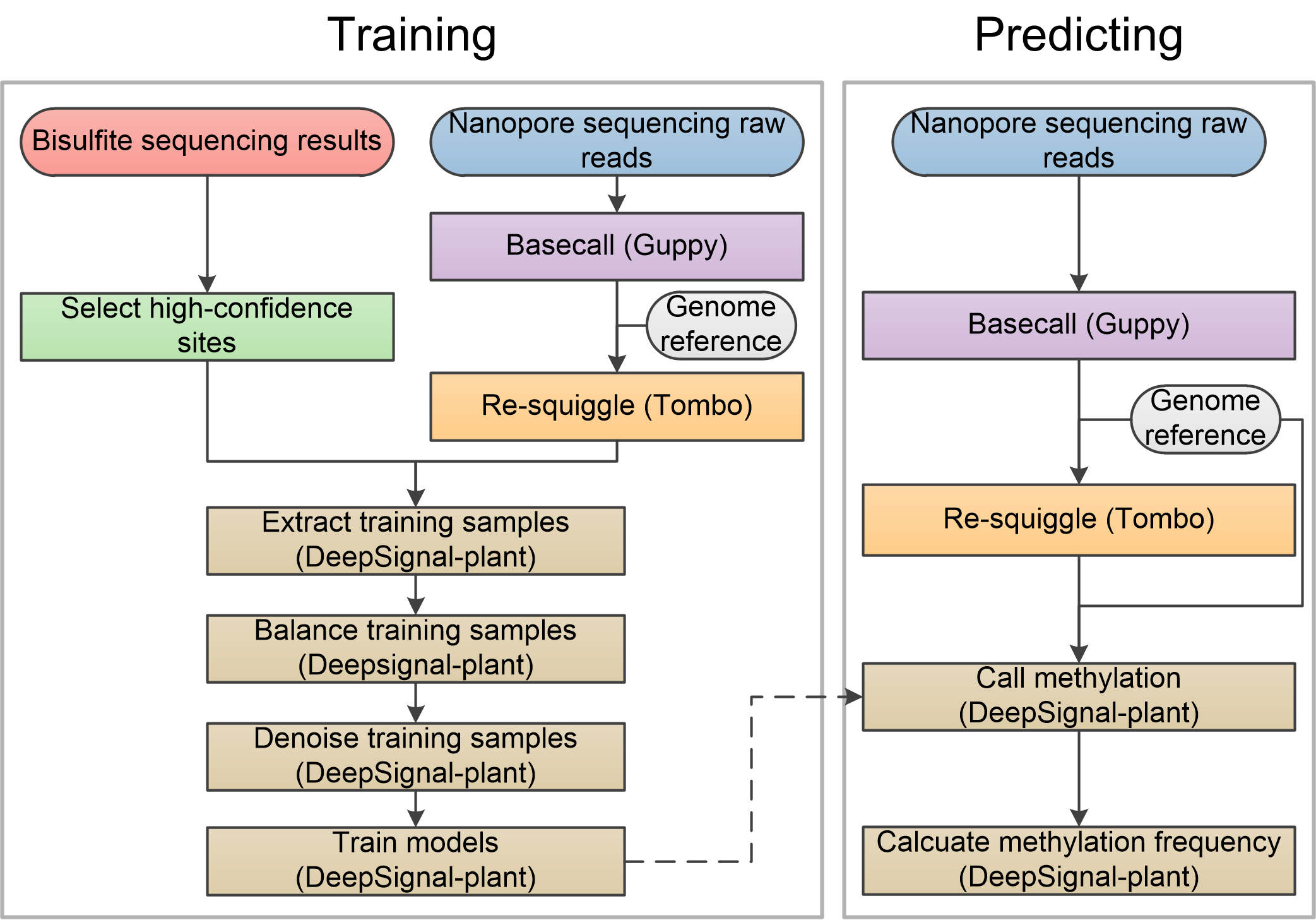

**Supplementary Fig. 4** Flowchart of our proposed pipeline.

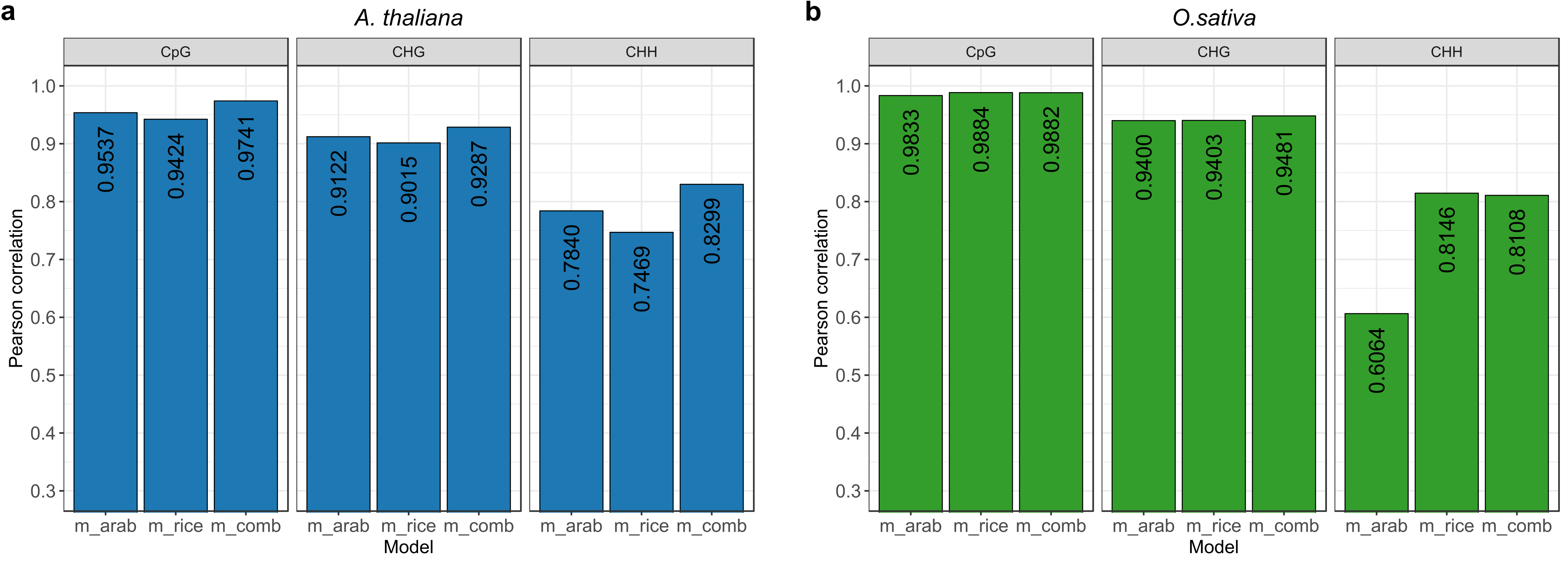

**Supplementary Fig. 5** 5mC detection in *A. thaliana* (**a**) and *O. sativa* (**b**) using models of DeepSignal-plant trained from different datasets. m_arab, m_rice, m_comb represent the models of DeepSignal-plant trained using ~500× *A. thaliana* Nanopore reads, ~115× *O. sativa* (sample1) Nanopore reads and the combined Nanopore reads, respectively. Pearson correlations are calculated using the results from ~20× Nanopore reads of *A.thaliana* and *O. sativa* (sample1) with the corresponding bisulfite replicates, respectively.

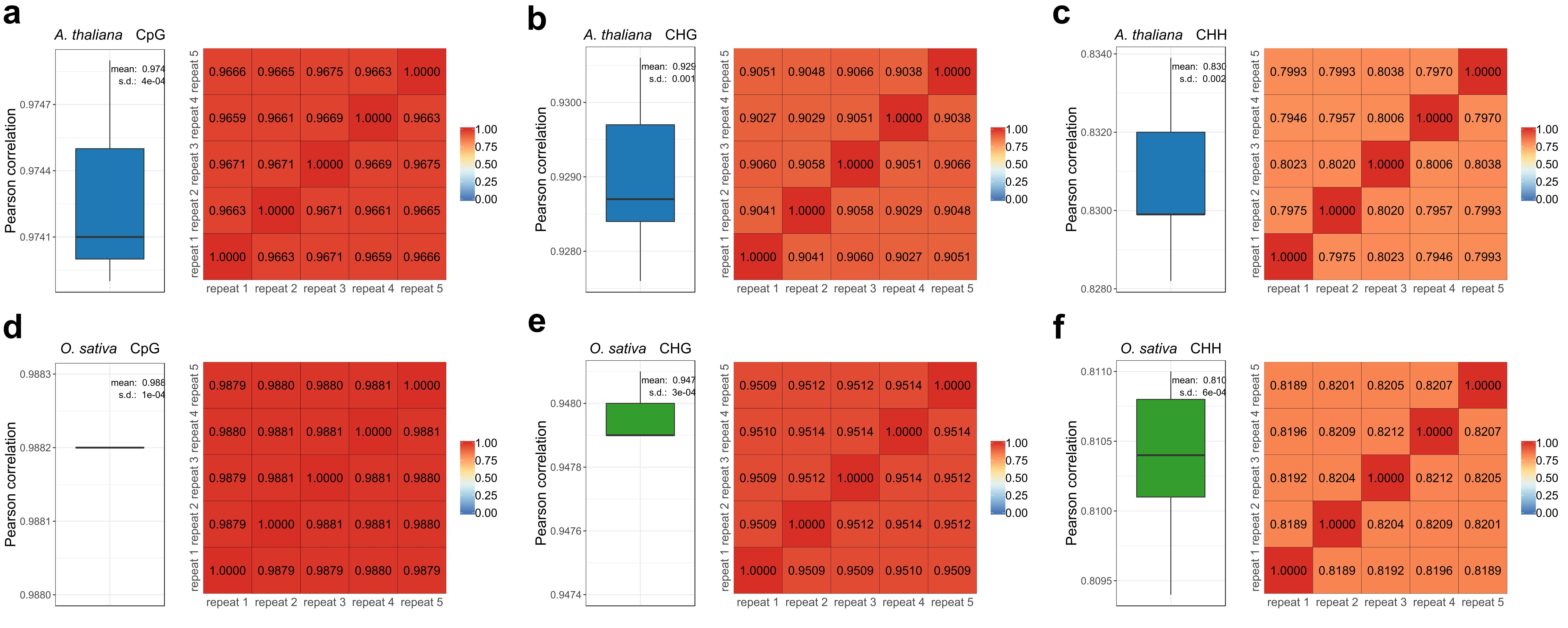

**Supplementary Fig. 6** Evaluation of our proposed pipeline by randomly selecting ~20× reads of *A. thaliana* and *O. sativa* (sample1) for 5 repeated times. **a-c:** CpG (a), CHG (b) and CHH (c) methylation of *A. thaliana*. **d-f:** CpG (a), CHG (b) and CHH (c) methylation of *O. sativa* (rep1). Boxplot: Pearson correlation with the results of bisulfite sequencing. Heatmap: Pearson correlation between the results of the 5 repeated tests. Models of DeepSignal-plant were trained by using combined reads of *A. thaliana* and *O. sativa*.

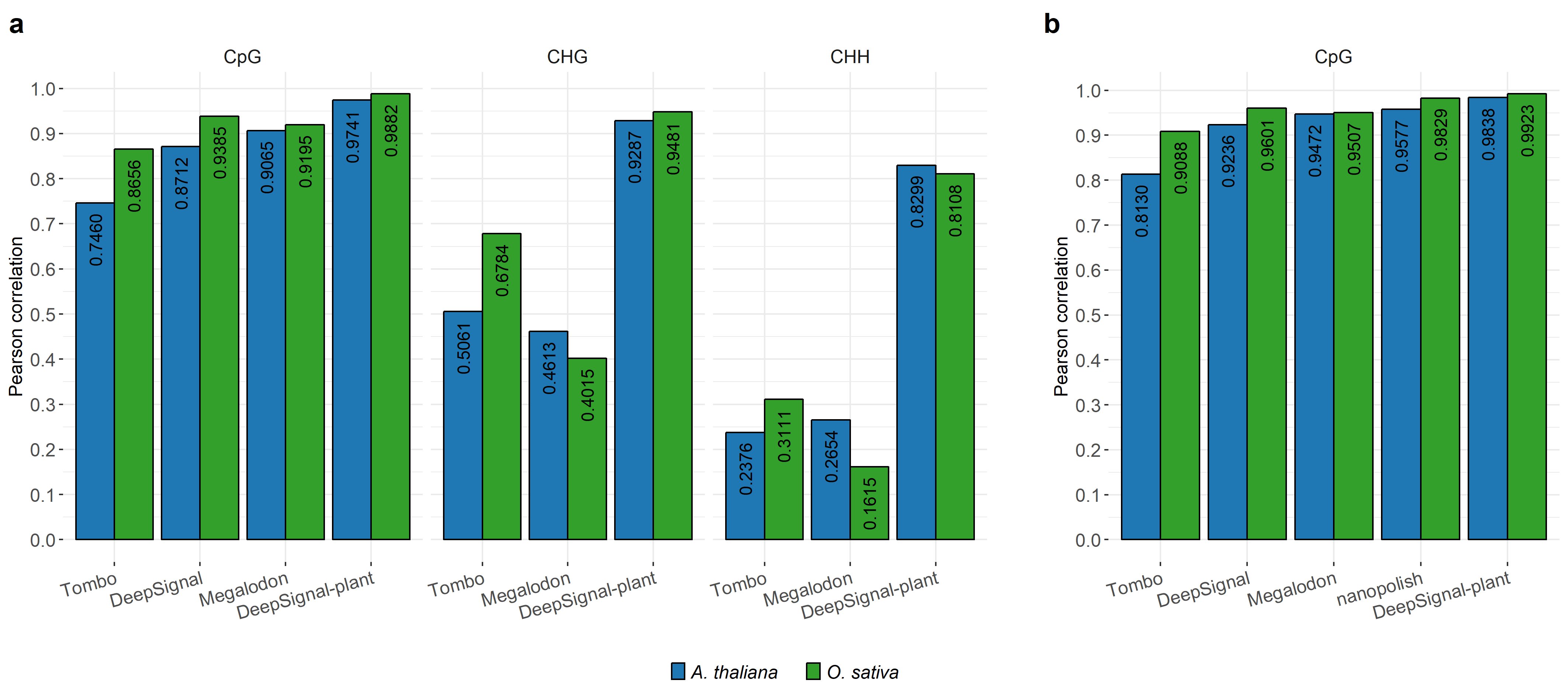

**Supplementary Fig. 7** Comparison of different methods for detecting methylation states of strand-specific sites (**a**) and non-strand-specific CpG sites (**b**) in *A. thaliana* and *O. sativa* (sample1). Pearson correlations are calculated using the results from ~20× Nanopore reads of *A. thaliana* and *O. sativa* (sample1) with the corresponding bisulfite replicates, respectively. Models of DeepSignal-plant were trained by using combined reads of *A. thaliana* and *O. sativa*.

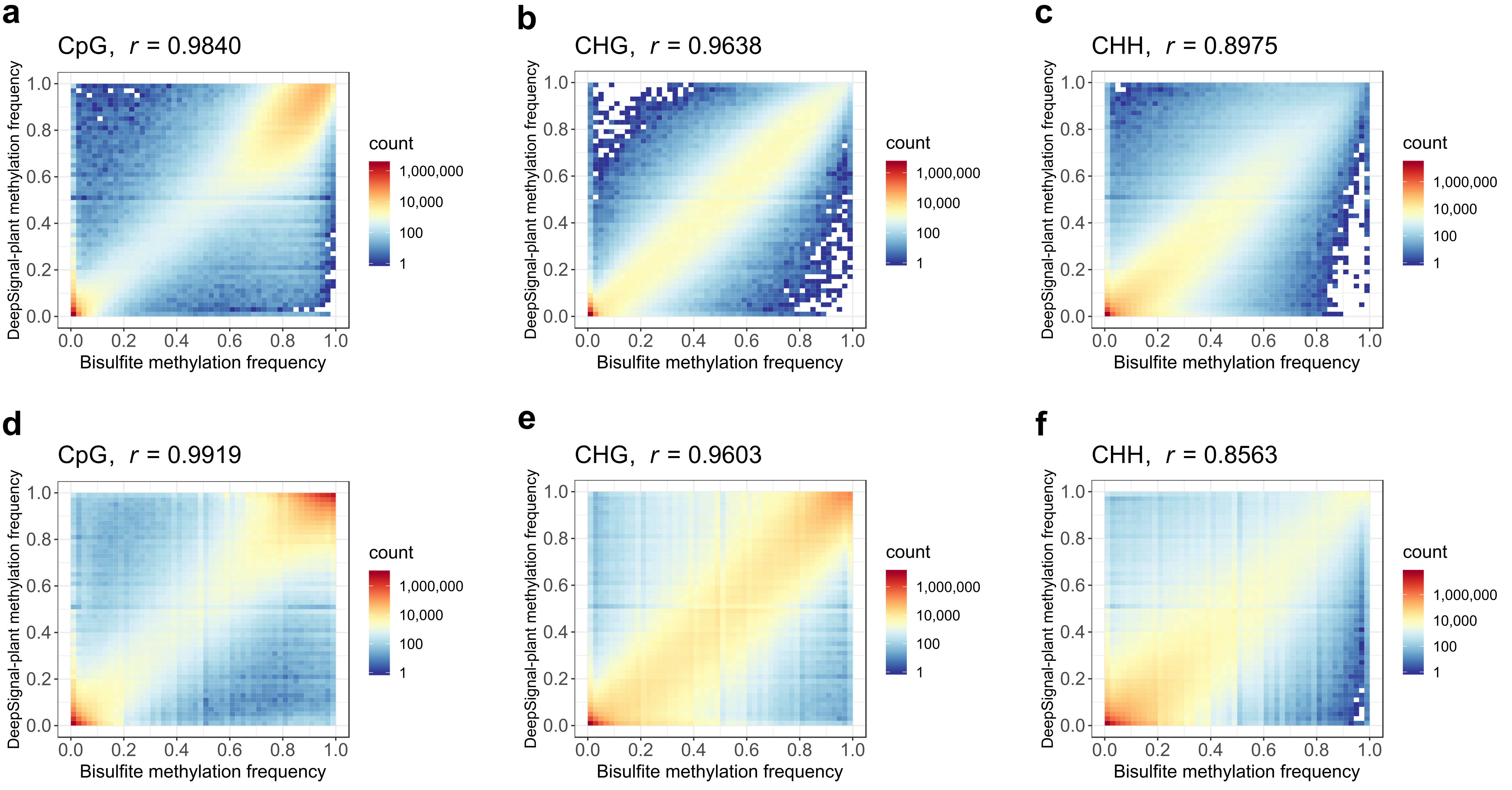

**Supplementary Fig. 8** Comparison of methylation frequencies of cytosines calculated by DeepSignal-plant and bisulfite sequencing. **a-c:** CpG (a), CHG (b), and CHH (c) methylation in *A. thaliana*. **d-f:** CpG (d), CHG (e), and CHH (f) methylation in *O. sativa* (sample1). *r* is Pearson correlation. ~100× coverage of Nanopore reads were used for *A. thaliana* and *O. sativa* (sample1), respectively. Models of Megalodon were trained by using combined reads of *A. thaliana* and *O. sativa*.

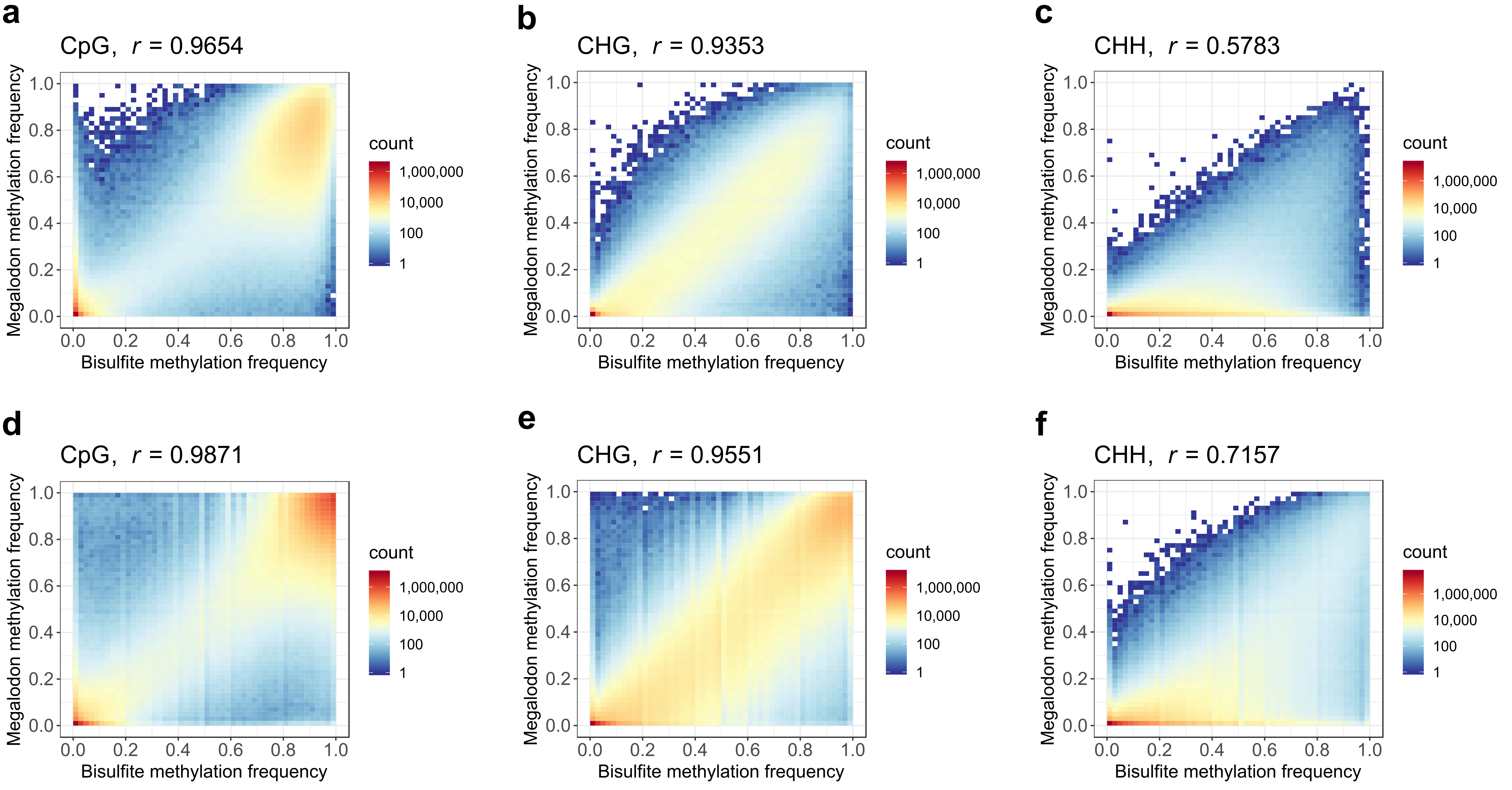

**Supplementary Fig. 9** Comparison of methylation frequencies of cytosines calculated by Megalodon and bisulfite sequencing. **a-c:** CpG (a), CHG (b), and CHH (c) methylation in *A. thaliana*. **d-f:** CpG (d), CHG (e), and CHH (f) methylation in *O. sativa* (sample1). *r* is Pearson correlation. ~100× coverage of Nanopore reads were used for *A. thaliana* and *O. sativa* (sample1), respectively. Models of Megalodon were trained by using combined reads of *A. thaliana* and *O. sativa*.

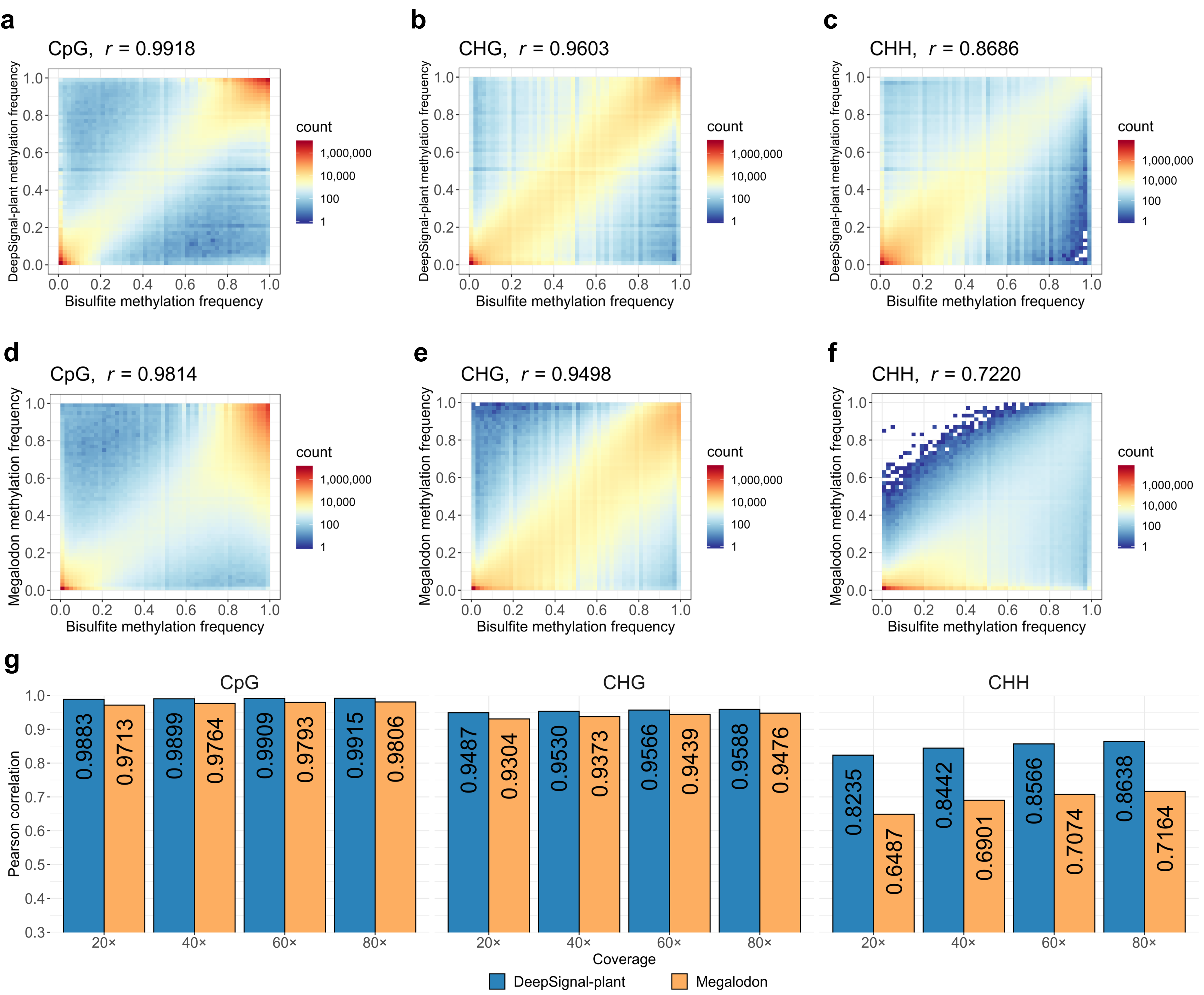

**Supplementary Fig. 10** Comparison of methylation frequencies of cytosines calculated by Nanopore sequencing (DeepSignal-plant and Megalodon) and bisulfite sequencing in *O. sativa* (sample2). **a-c:** Comparison of DeepSignal-plant and bisulfite sequencing. **d-f:** Comparison of Megalodon and bisulfite sequencing. *r* is Pearson correlation. ~100× coverage of Nanopore reads of *O. sativa* (sample2) were used. **g:** Comparison between DeepSignal-plant and Megalodon on 5mC detection against bisulfite sequencing in CpG, CHG and CHH contexts under different coverage of Nanopore reads. For each coverage (20× to 80×), the reads were randomly shuffled and selected from ~100× reads. Values for each coverage are average of 5 replicated tests. Models of Megalodon and DeepSignal-plant were trained by using combined reads of *A. thaliana* and *O. sativa*.

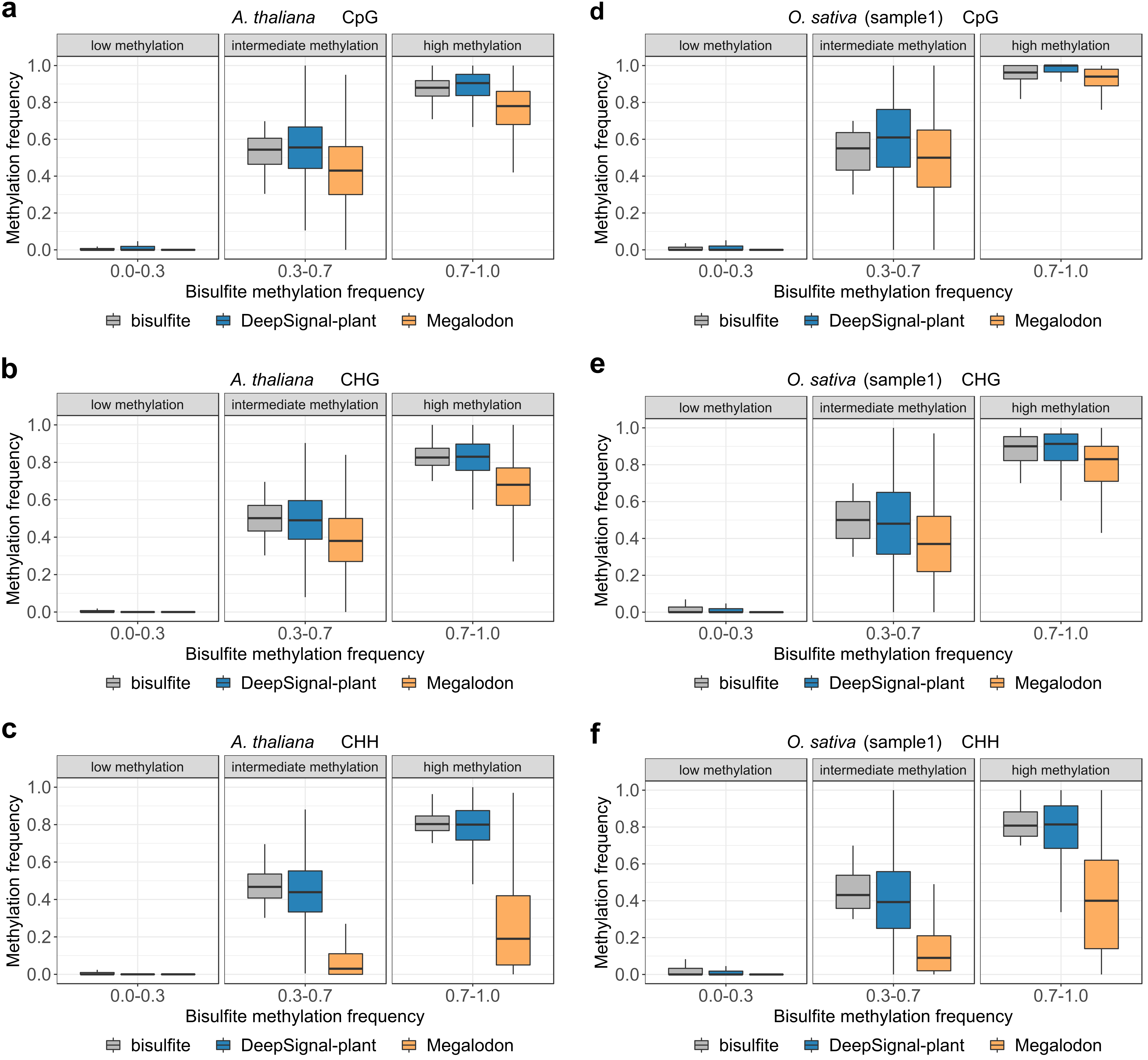

**Supplementary Fig. 11** Distribution of methylation frequencies called by DeepSignal-plant and Megalodon from ~100× Nanopore reads of *A. thaliana* and *O. sativa* (sample1) against bisulfite sequencing across three methylation bins: low methylation (0.0-0.3), intermediate methylation (0.3-0.7), and high methylation (0.7-1.0). **a-c**: CpG (a), CHG (b), and CHH (c) methylation in *A. thaliana*. **d-f**: CpG (d), CHG (e), and CHH (f) methylation in *O. sativa* (sample1). Models of Megalodon and DeepSignal-plant were trained by using combined reads of *A. thaliana* and *O. sativa*.

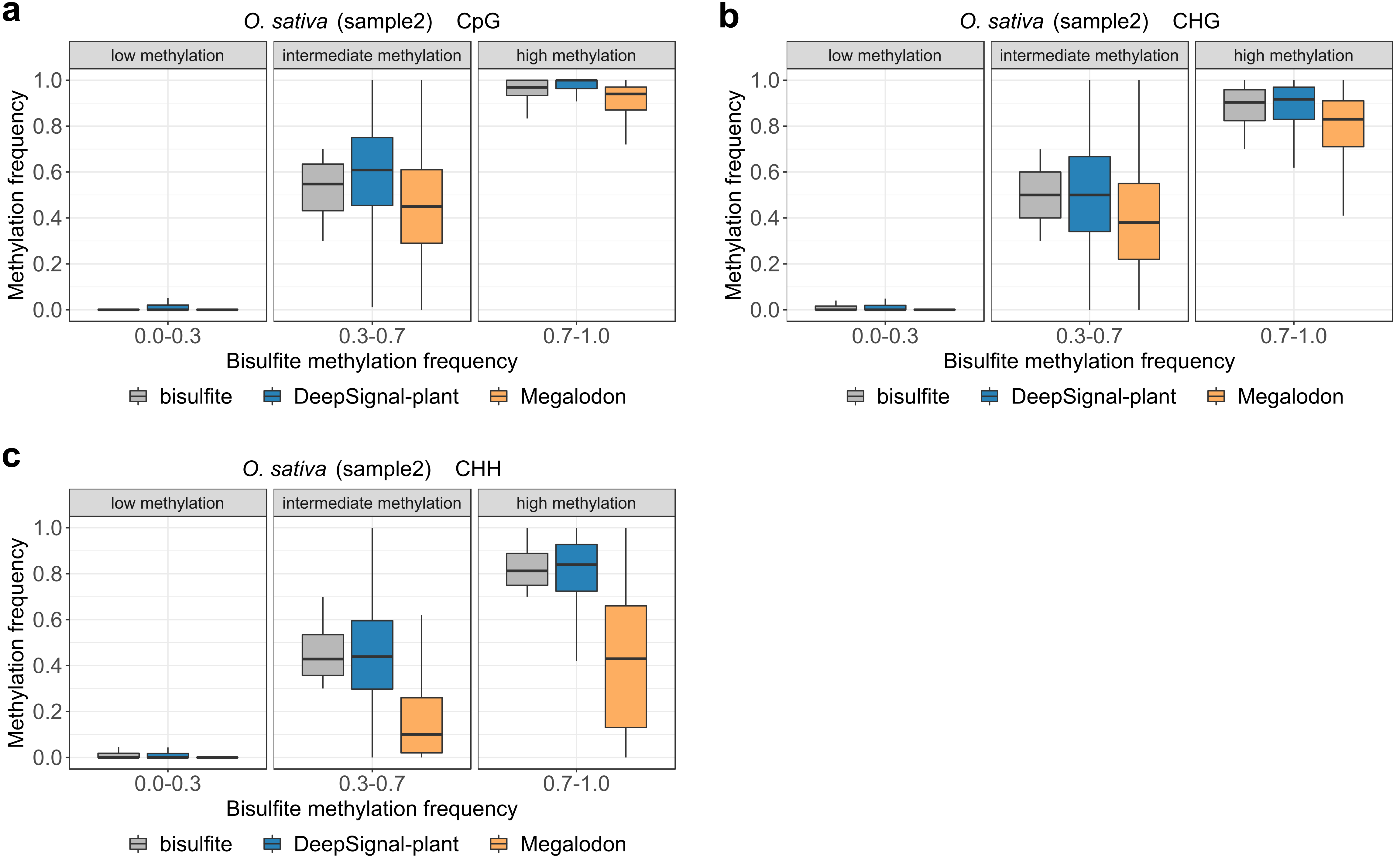

**Supplementary Fig. 12** Distribution of methylation frequencies called by DeepSignal-plant and Megalodon from ~100× Nanopore reads of *O. sativa* (sample2) against bisulfite sequencing across three methylation bins: low methylation (0.0-0.3), intermediate methylation (0.3-0.7), and high methylation (0.7-1.0). **a-c**: CpG (a), CHG (b), and CHH (c) methylation in *O. sativa* (sample2). Models of DeepSignal-plant and Megalodon were trained by using combined reads of *A. thaliana* and *O. sativa*.

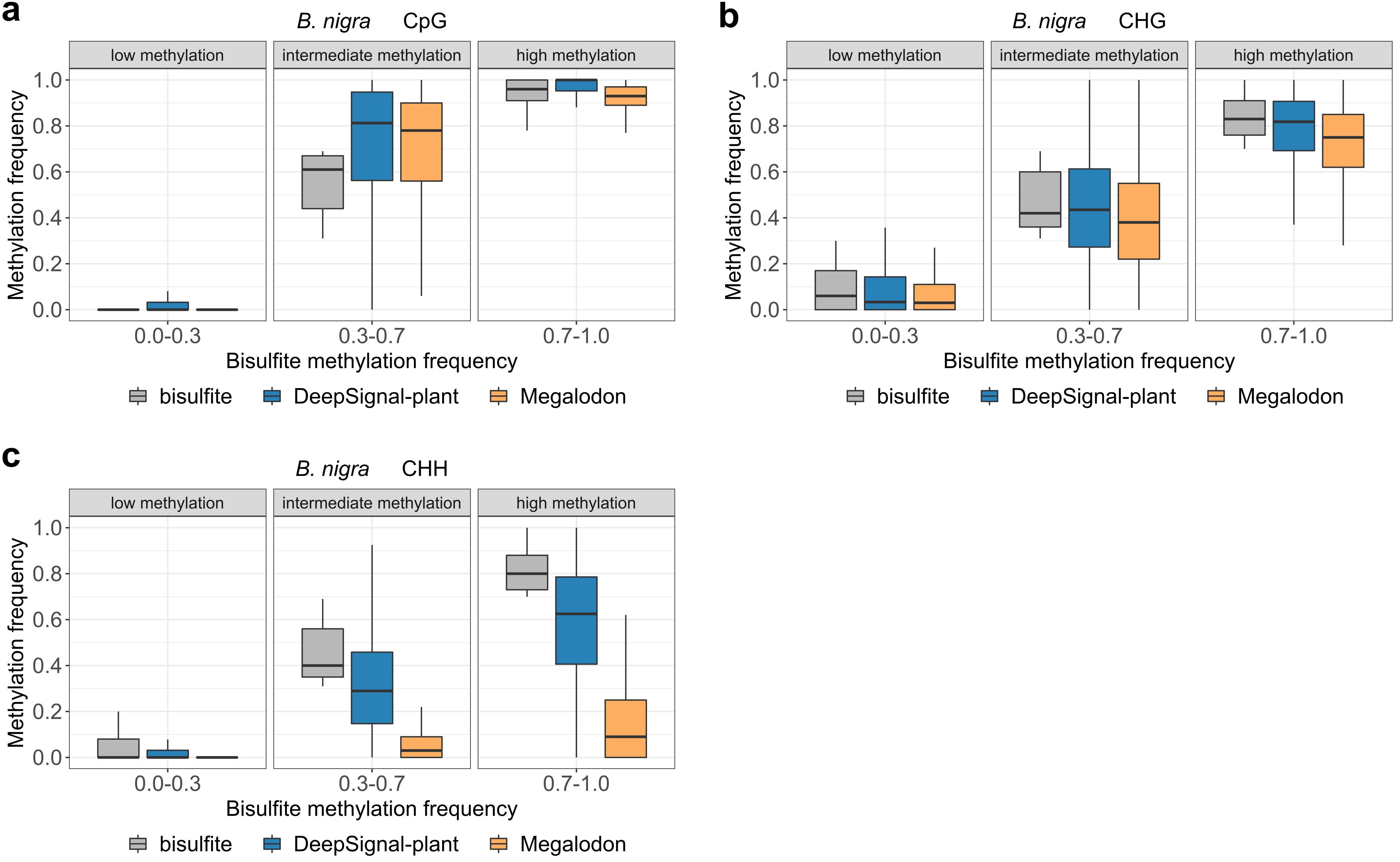

**Supplementary Fig. 13** Distribution of methylation frequencies predicted by DeepSignal-plant and Megalodon from ~78× Nanopore reads of *B. nigra* against bisulfite sequencing across three methylation bins: low methylation (0.0-0.3), intermediate methylation (0.3-0.7), and high methylation (0.7-1.0). **a**: CpG motif. **b**: CHG motif. **c**: CHH motif. Models of DeepSignal-plant and Megalodon were trained by using combined reads of *A. thaliana* and *O. sativa*.

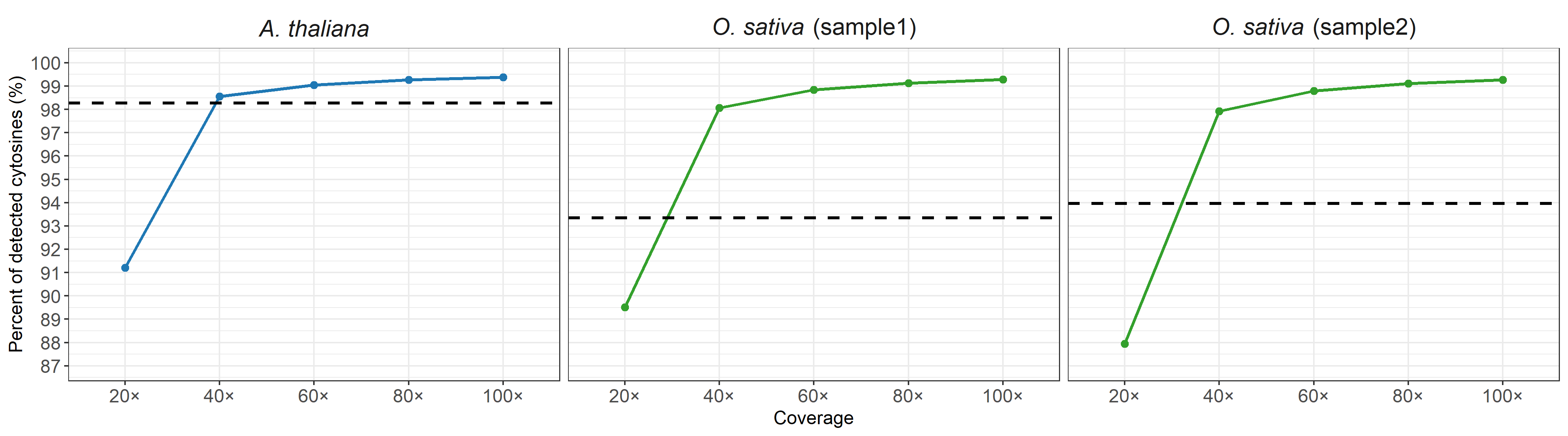

**Supplementary Fig. 14** Percent of cytosines detected by DeepSignal-plant from Nanopore sequencing (coverage>=5) in genomes of *A. thaliana* and *O. sativa*. Reads for coverage 20× to 80× are randomly shuffled and selected from ~100× reads. Values for coverage 20× to 80× are average of 5 replicated tests. Black dash lines indicate percent of cytosines detected by bisulfite sequencing (coverage>=5).

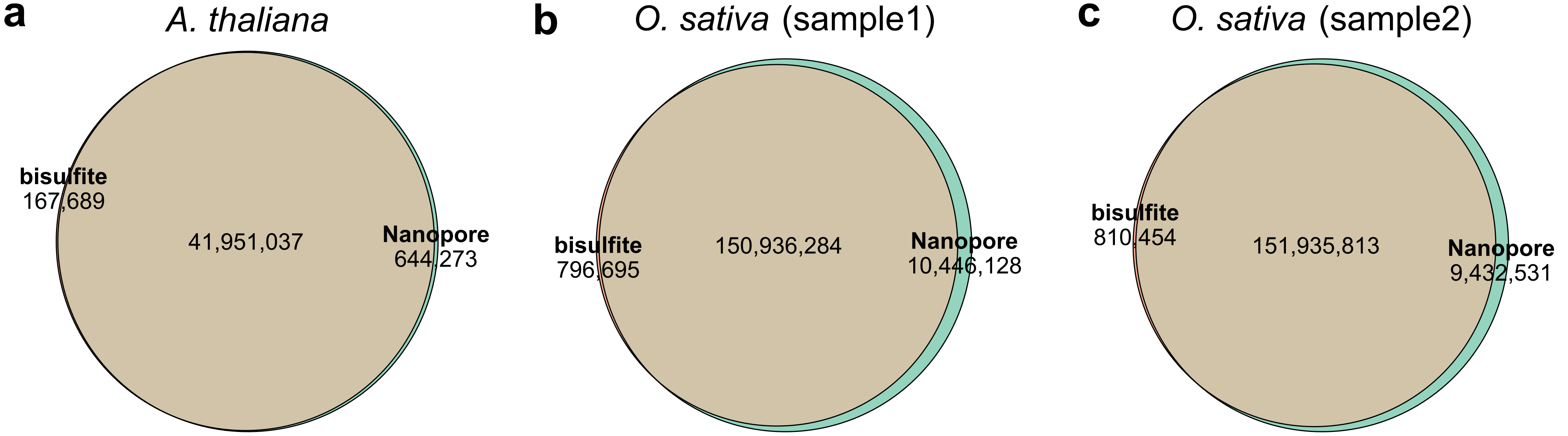

**Supplementary Fig. 15** Comparison of cytosines detected by bisulfite sequencing (Bismark) and Nanopore sequencing (DeepSignal-plant, ~100×). **a:** A. thaliana. **b:** *O. sativa* (sample1). **c:** *O. sativa* (sample2)

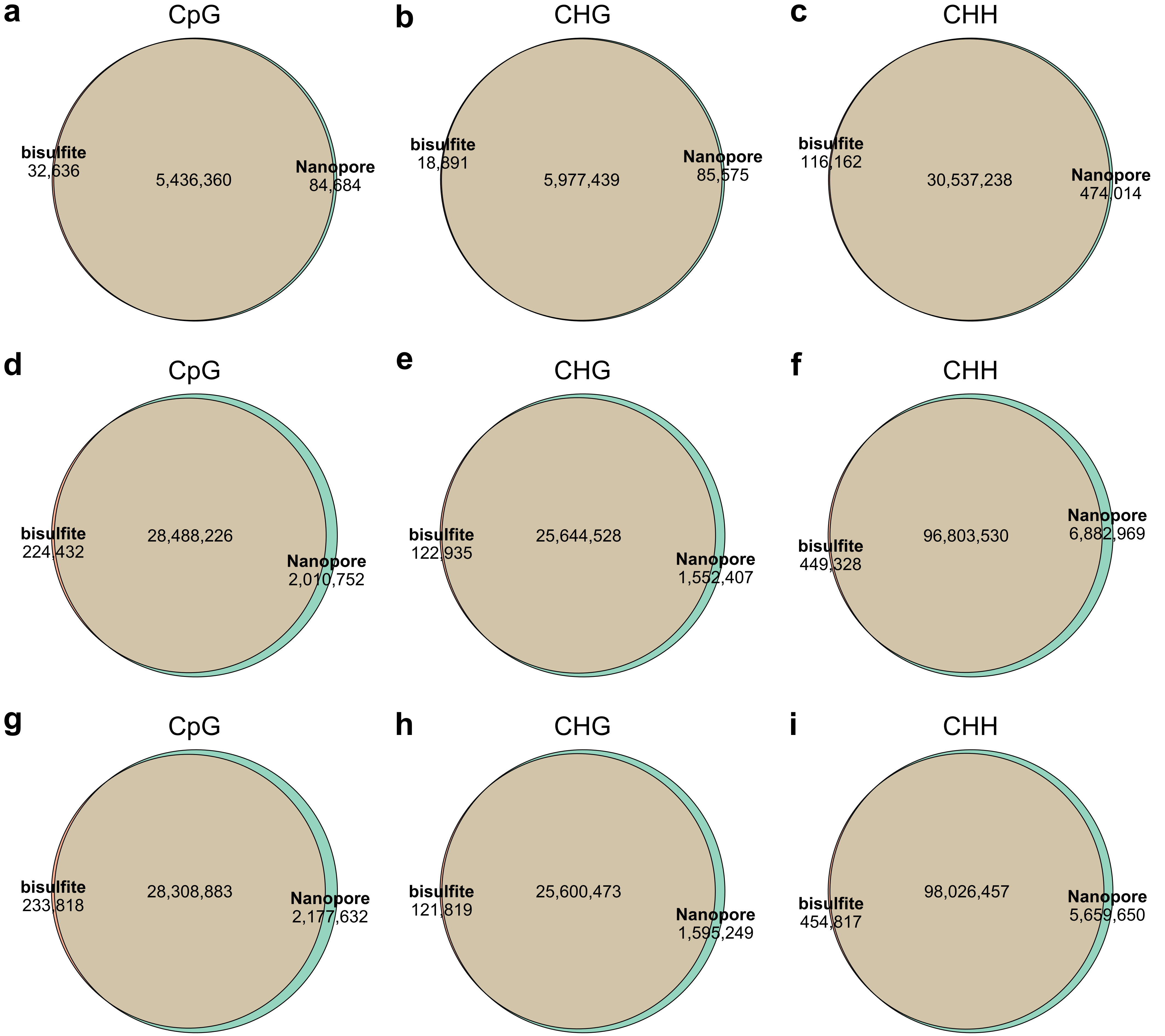

**Supplementary Fig. 16** Comparison of cytosines detected by bisulfite sequencing (Bismark) and Nanopore sequencing (DeepSignal-plant, ~100×) in three motifs. **a-c**: Comparison of number of CpG (a), CHG (b) and CHH (c) sites in *A. thaliana*. **d-f:** Comparison of number of CpG (d), CHG (e) and CHH (f) sites in *O. sativa* (sample1). **g-i:** Comparison of number of CpG (g), CHG (h) and CHH (i) sites in *O. sativa* (sample2).

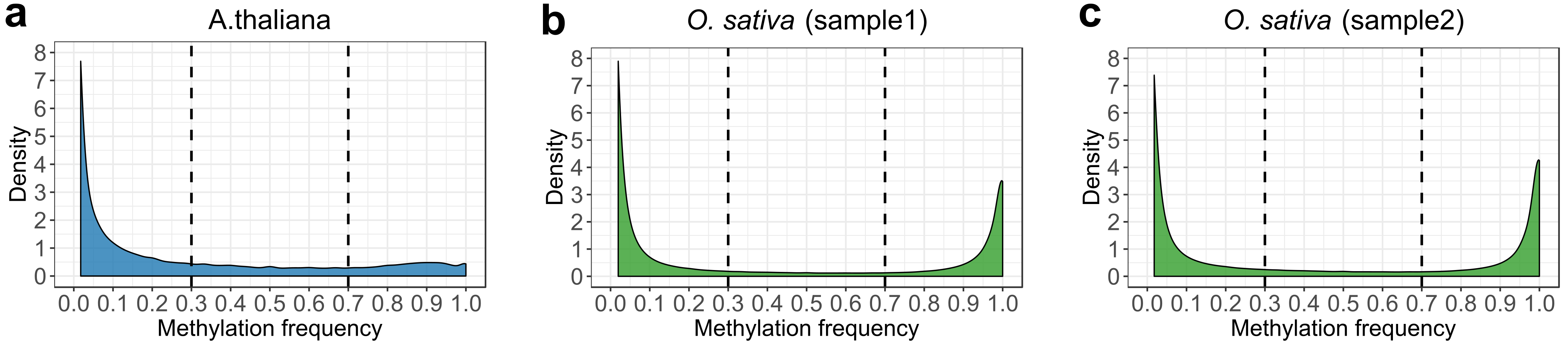

**Supplementary Fig. 17** Methylation frequencies of cytosines which can only be detected by Nanopore sequencing (DeepSignal-plant) in *A. thaliana* (**a**), *O. sativa* sample1 (**b**) and sample2 (**c**).

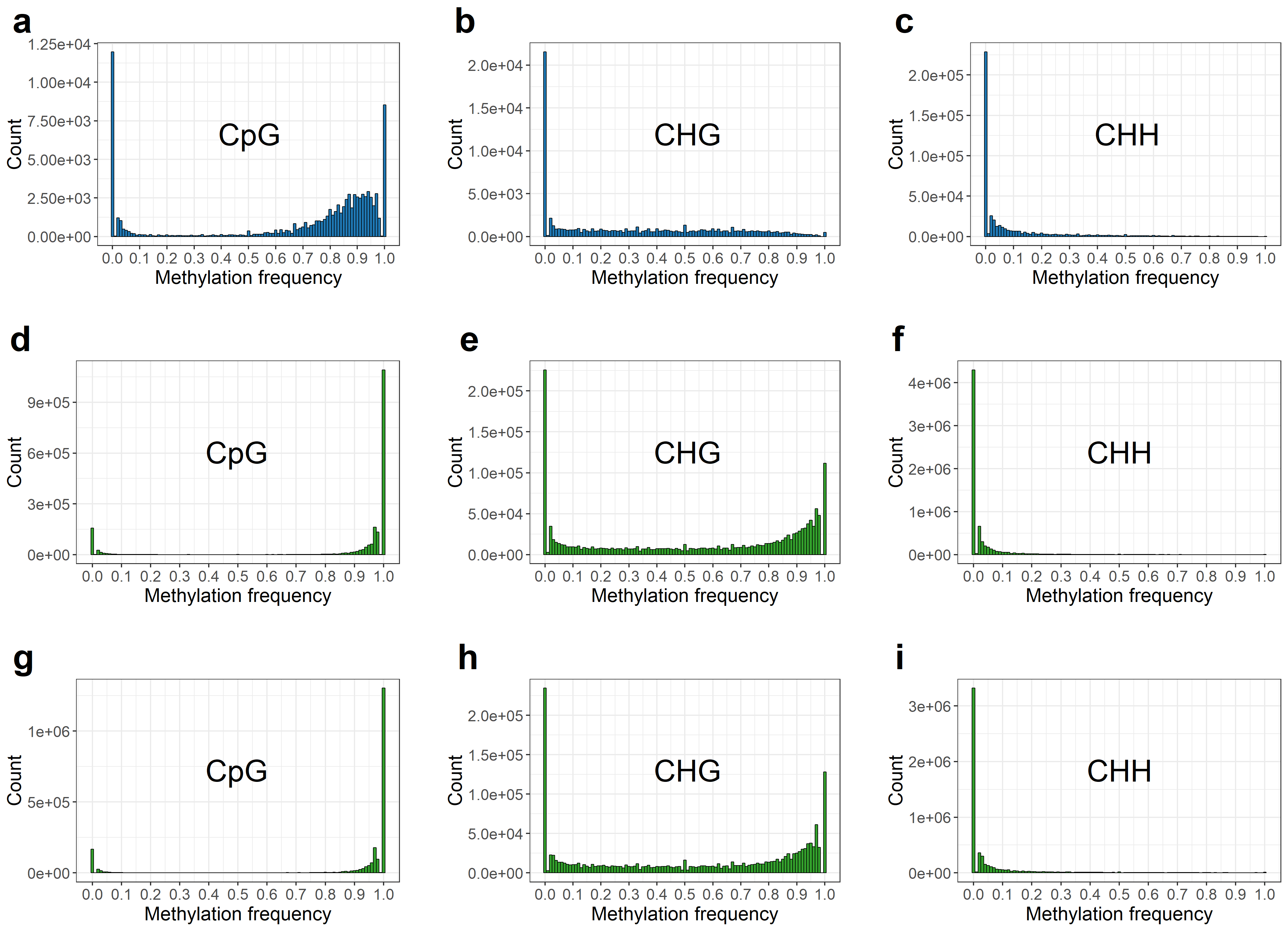

**Supplementary Fig. 18** Methylation frequencies of cytosines which can only be detected by Nanopore sequencing (DeepSignal-plant). **a-c:** Methylation frequencies of CpG (a), CHG (b) and CHH (c) sites in *A. thaliana*. **d-f:** Methylation frequencies of CpG (d), CHG (e) and CHH (f) sites in *O. sativa* (sample1). **g-i:** Methylation frequencies of CpG (g), CHG (h) and CHH (i) sites in *O. sativa* (sample2).

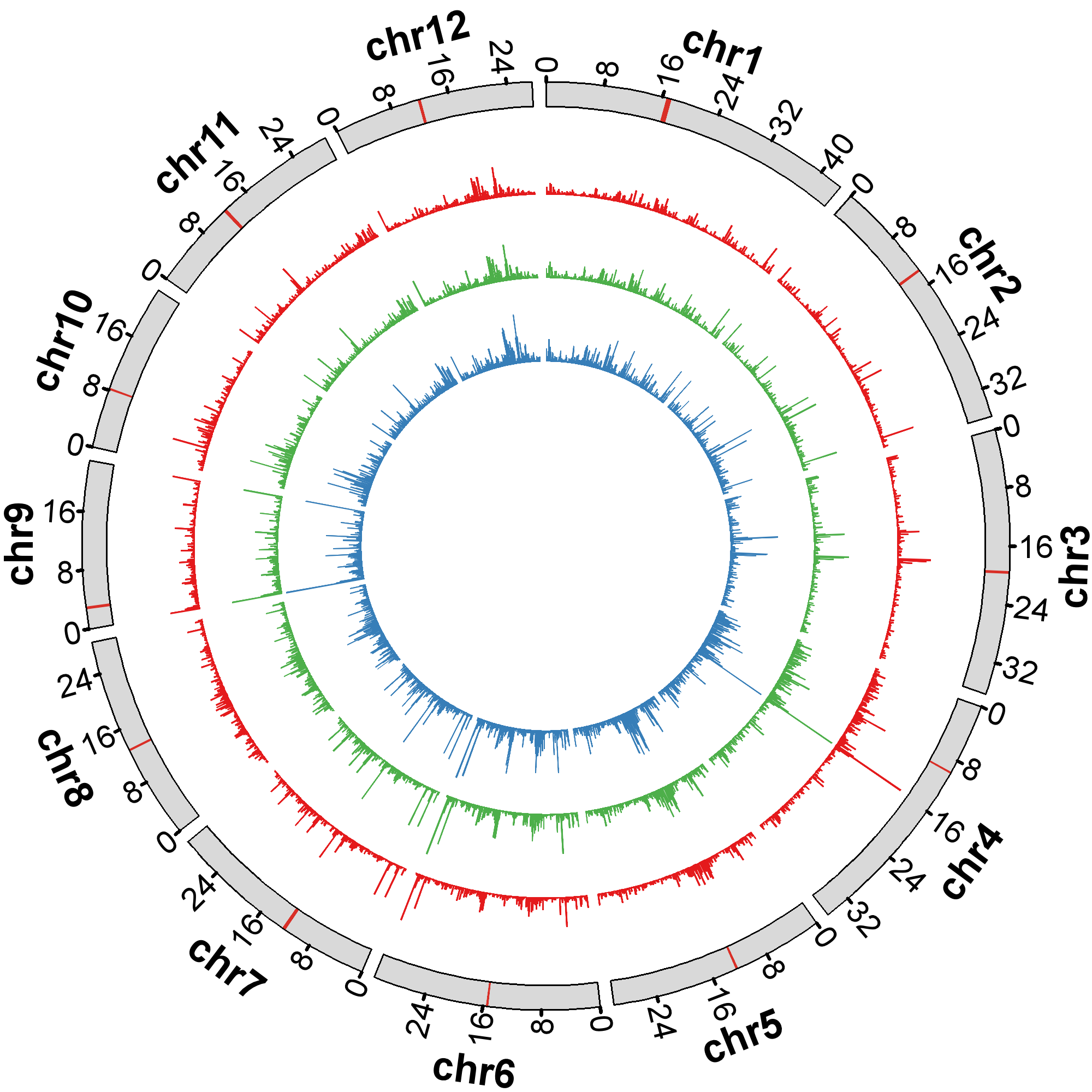

**Supplementary Fig. 19** Circos plot of number of cytosines detected by Nanopore sequencing only in the *O. sativa* (sample2). Cycles from inner to outer: CpG (blue), CHG (green), CHH (red), reference (the chromosomes are binned into 200,000-bp (base pair) windows. The centromeric region is indicted by red bar in each chromosome).

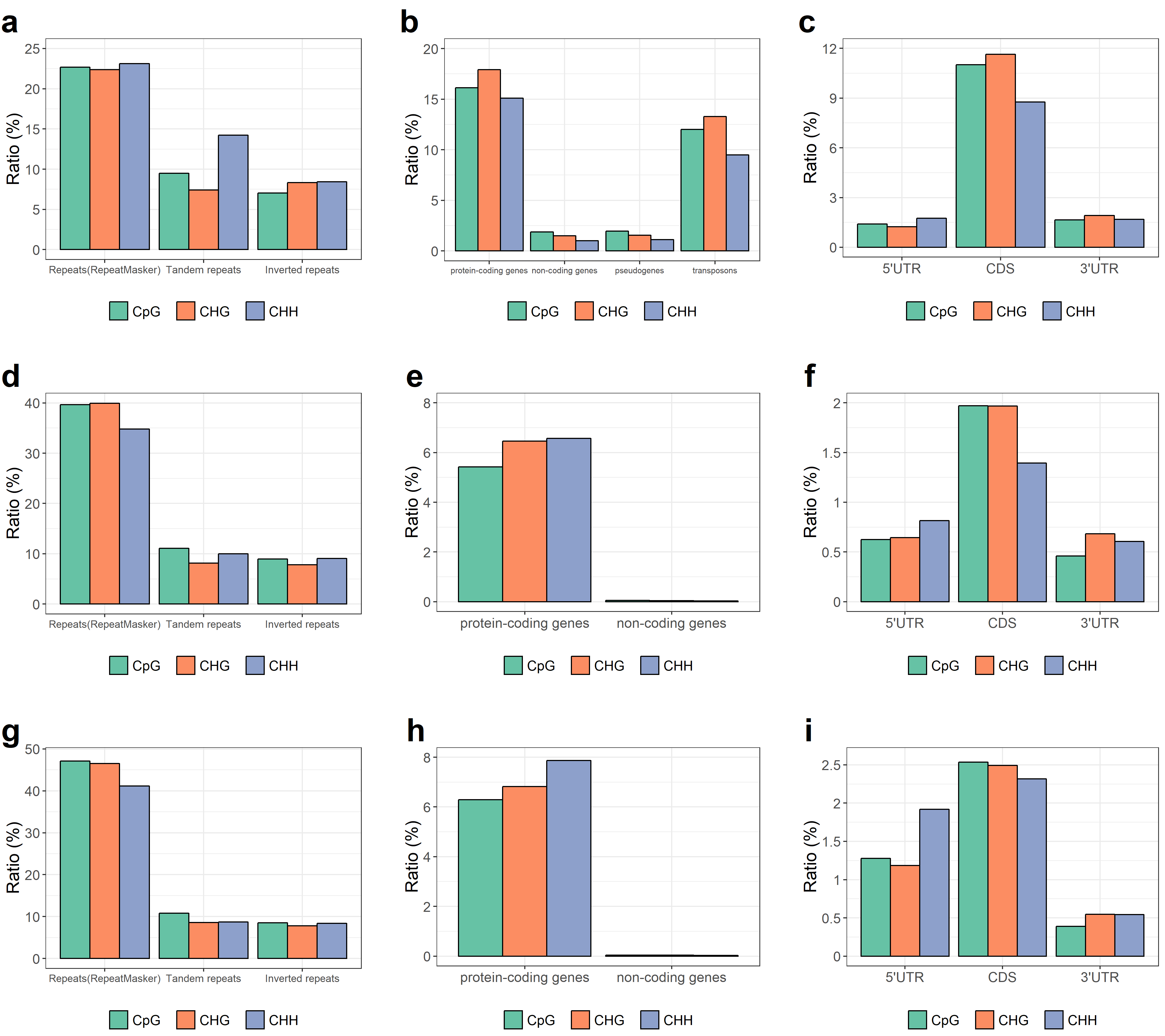

**Supplementary Fig. 20** Distribution of cytosines which can only be detected by Nanopore sequencing (DeepSignal-plant) in repeats and gene regions. **a-c:** Proportion of cytosines which can only be detected by Nanopore sequencing in repeat regions (a), different kinds of genes (b) and gene bodies (c) of *A. thaliana*. **d-f:** Proportion of cytosines which can only be detected by Nanopore sequencing in repeat regions (d), different kinds of genes (e) and gene bodies (f) of *O. sativa* (sample1). **g-i:** Proportion of cytosines which can only be detected by Nanopore sequencing in repeat regions (g), different kinds of genes (h) and gene bodies (i) of *O. sativa* (sample2).

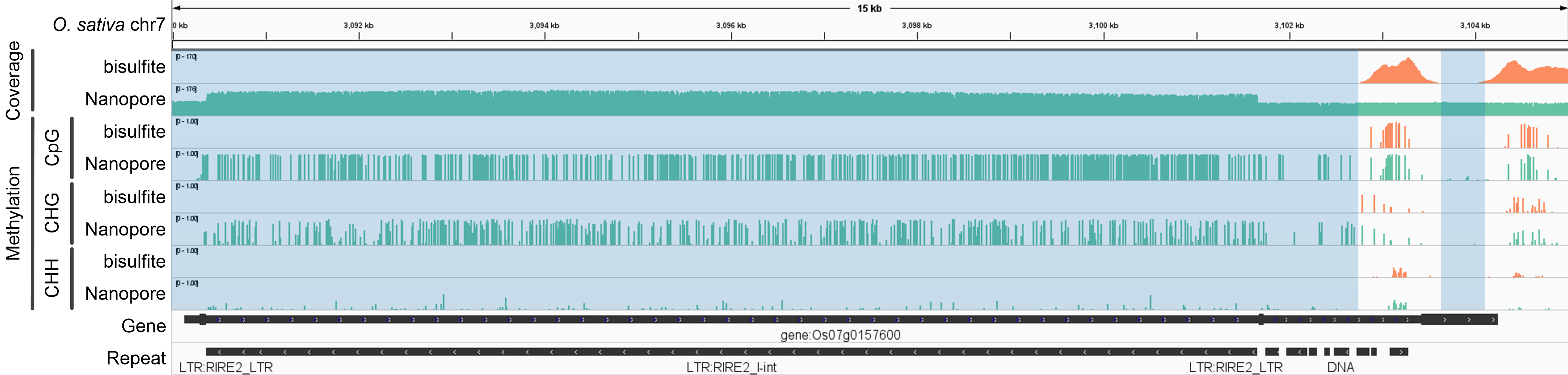

**Supplementary Fig. 21** Genome browser view [1] of the reads coverage and methylation in a 14 kb region of *O. sativa* (sample2) chr7 detected by bisulfite sequencing (Bismark) and Nanopore sequencing (DeepSignal-plant). The blue shaded area shows the gaps which cannot be mapped by bisulfite sequencing.

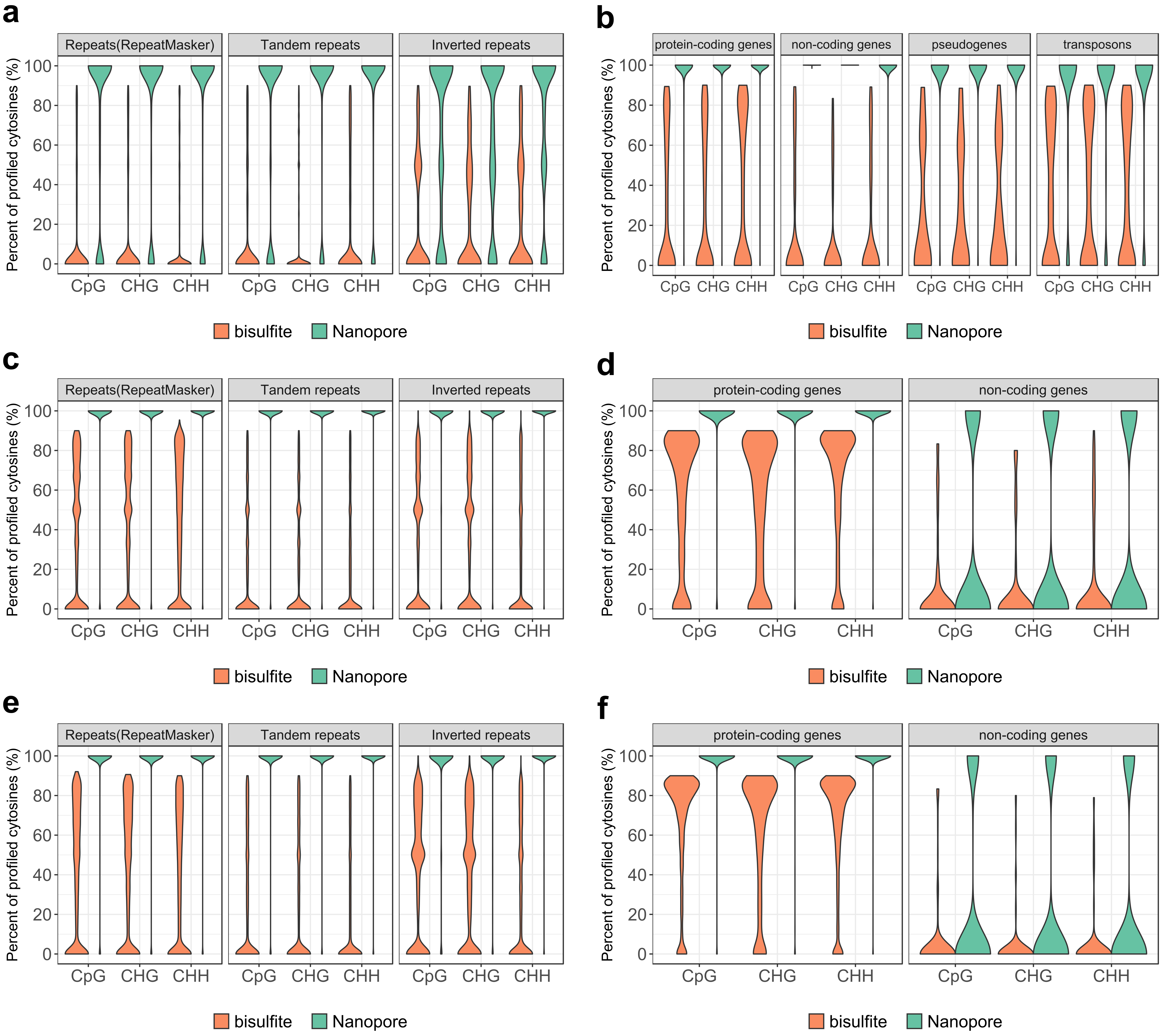

**Supplementary Fig. 22** Percent of profiled cytosines by bisulfite sequencing (Bismark) and Nanopore sequencing (DeepSignal-plant) in biological regions that cannot be fully profiled by bisulfite sequencing. Repeat regions and gene regions in which the percent of profiled cytosines by bisulfite sequencing <= 90% are selected for comparison. **a-b:** Comparison of repeat regions (a) and gene regions (b) in *A. thaliana*. **c-d:** Comparison of repeat regions (c) and gene regions (d) in *O. sativa* (sample1). **e-f:** Comparison of repeat regions (e) and gene regions (f) in *O. sativa* (sample2).

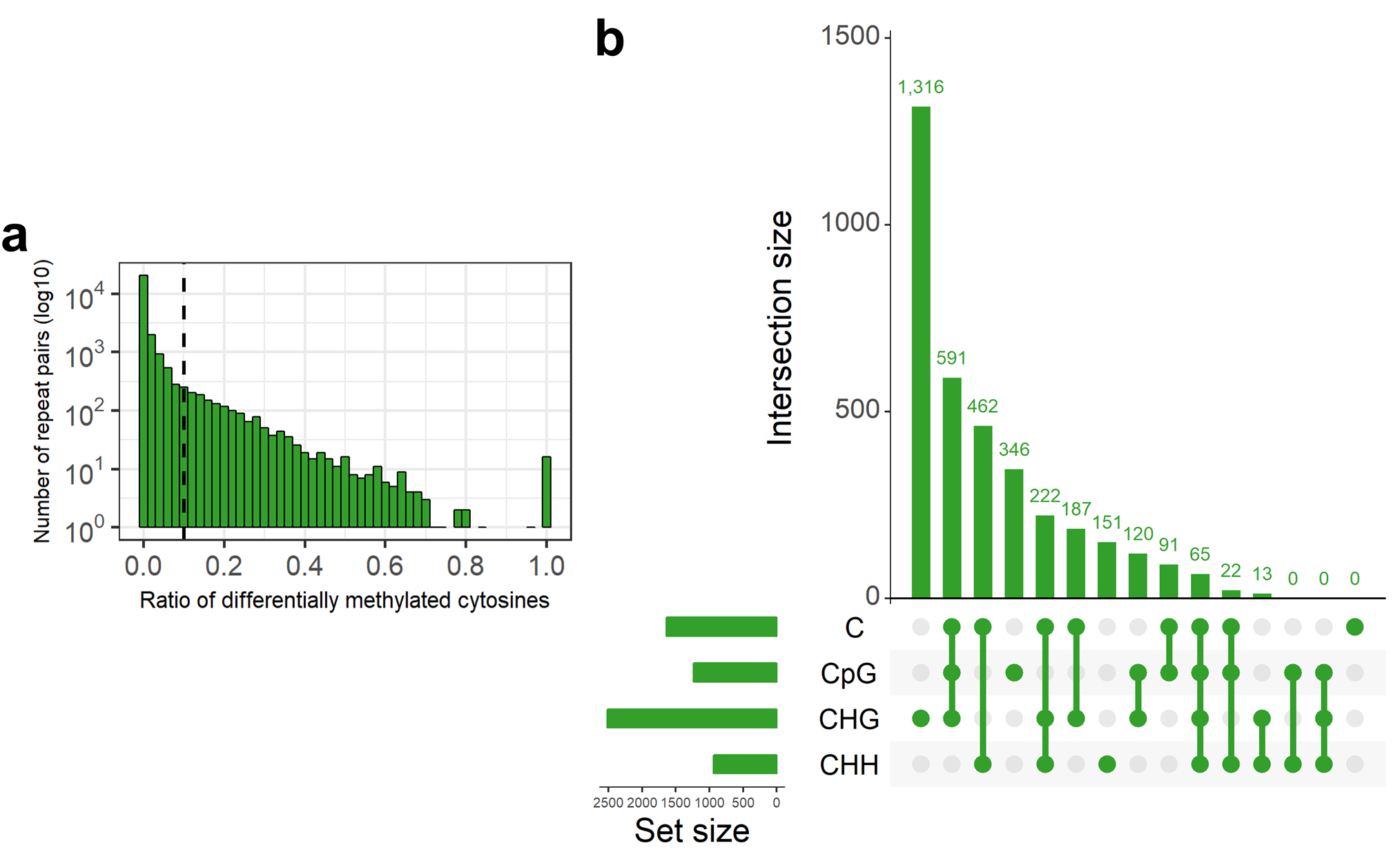

**Supplementary Fig. 23** Our proposed pipeline identified differentially methylated repeat pairs in *O. sativa* (rep2). **a:** Ratio of differentially methylated cytosines in repeat pairs. **c-d:** Matrix layout for all intersections of four sets of differentially methylated repeat pairs profiled by cytosines, CpG sites, CHG sites, and CHH sites independently. Circles in the matrix indicate sets that are part of the intersection.

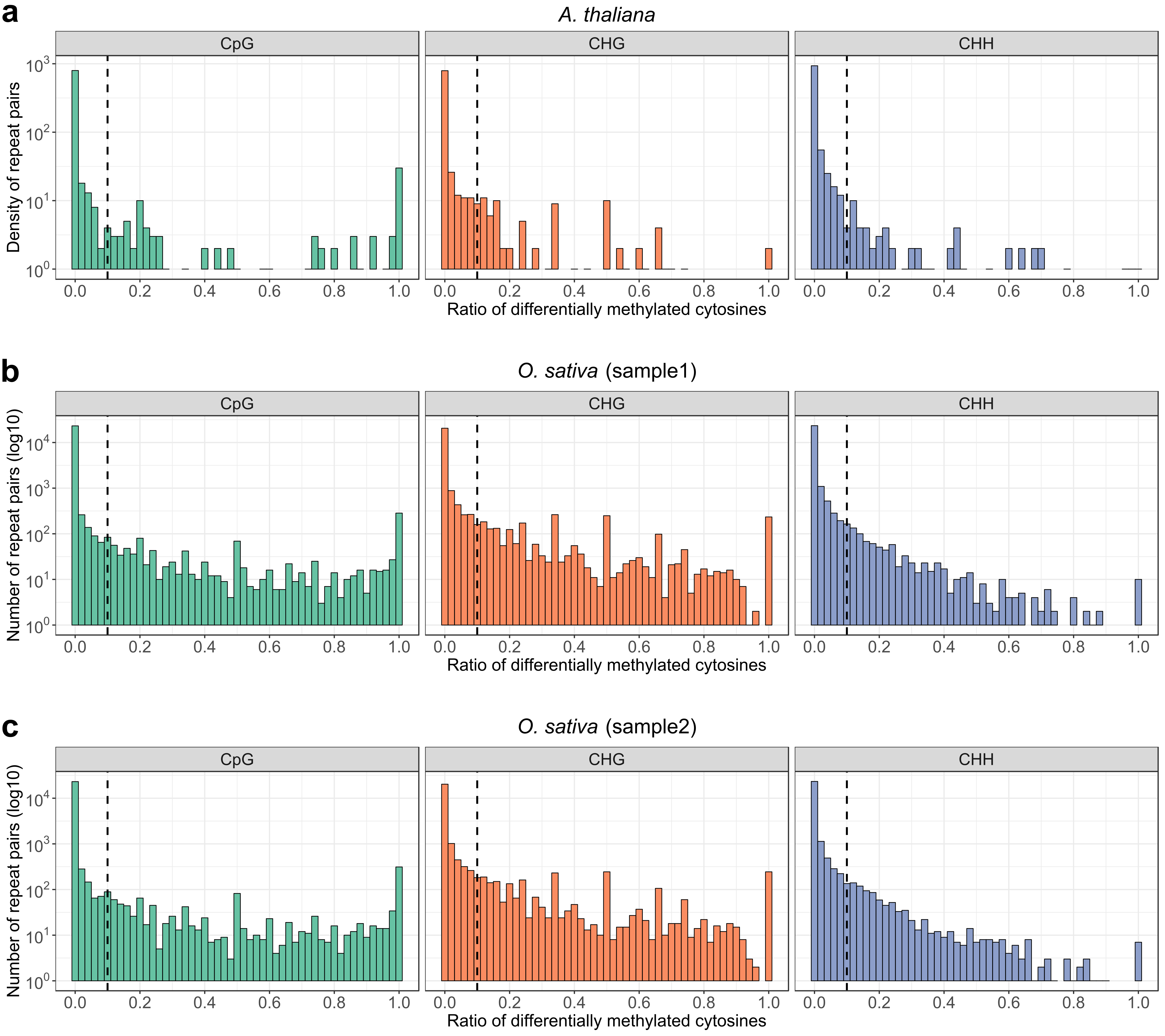

**Supplementary Fig. 24** Ratio of differentially methylated CpG, CHG, and CHH sites in repeat pairs of *A. thaliana* (a) and *O. sativa* sample1 (b) and *O. sativa* sample2 (c).

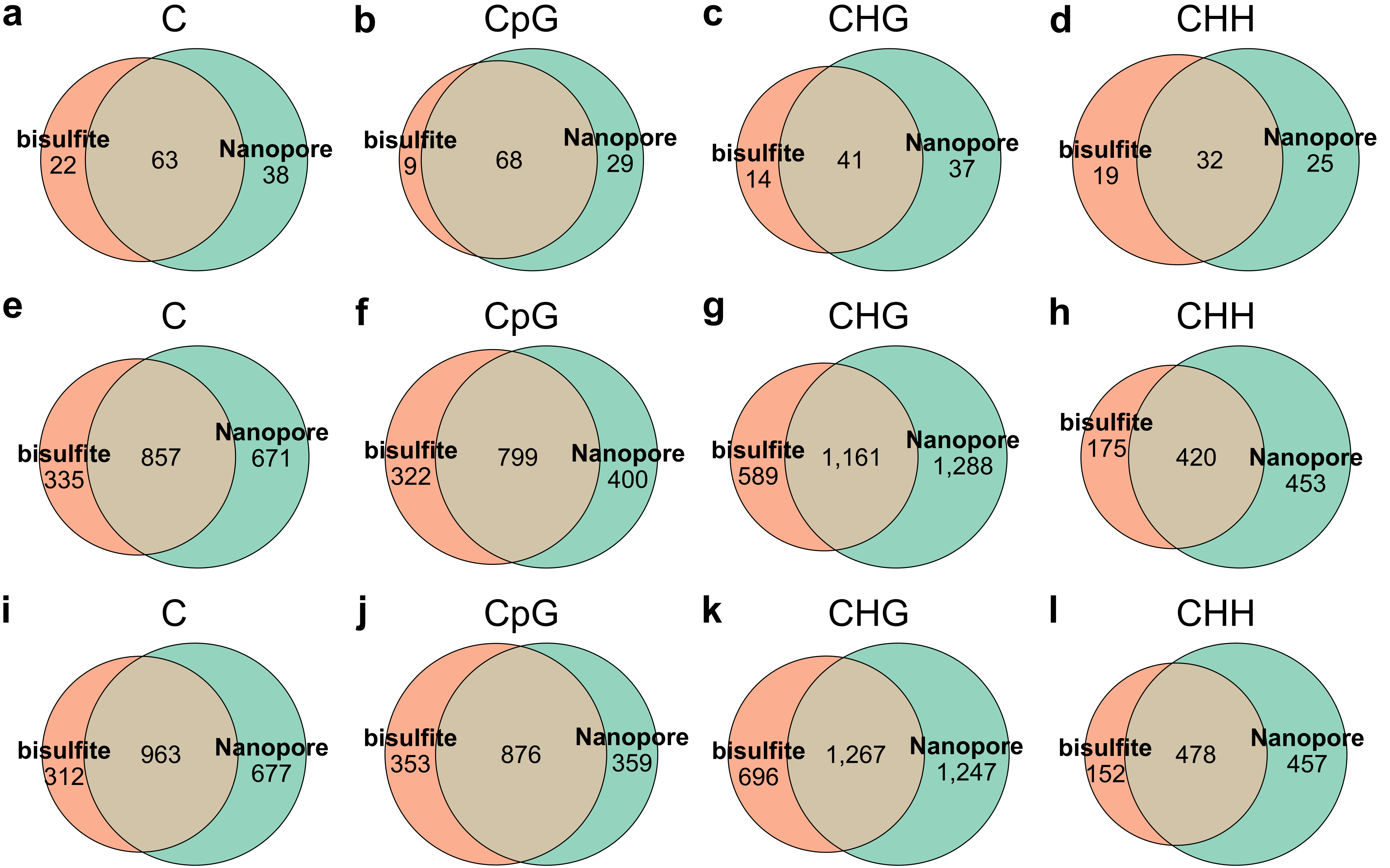

**Supplementary Fig. 25** Comparison of differentially methylated repeat pairs profiled by bisulfite sequencing (Bismark) and Nanopore sequencing (DeepSignal-plant). **a-d:** Comparison of differentially methylated repeat pairs identified by methylation of cytosines (a), CpG sites (b), CHG sites (c), and CHH sites (d) in *A. thaliana*. **e-h:** Comparison of differentially methylated repeat pairs identified by methylation of cytosines (e), CpG sites (f), CHG sites (g), and CHH sites (h) in *O. sativa* (sample1). **i-l:** Comparison of differentially methylated repeat pairs identified by methylation of cytosines (i), CpG sites (j), CHG sites (k), and CHH sites (l) in *O. sativa* (sample2).

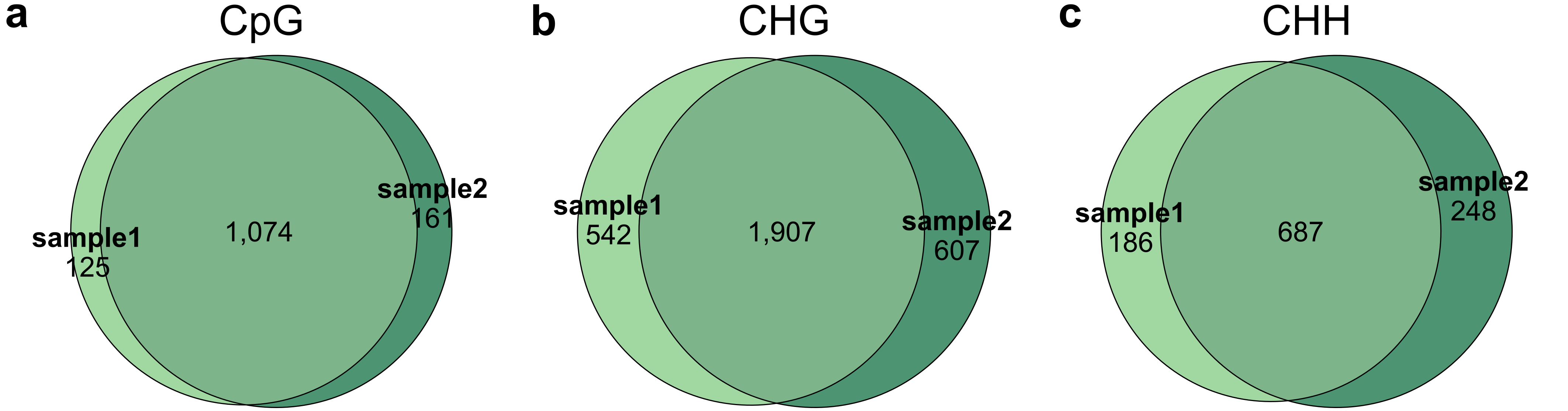

**Supplementary Fig. 26** Comparison of differentially methylated repeat pairs in *O. sativa* sample1 and sample2 identified by methylation of CpG sites (a), CHG sites (b), and CHH sites (c) independently, which were detected by DeepSignal-plant.

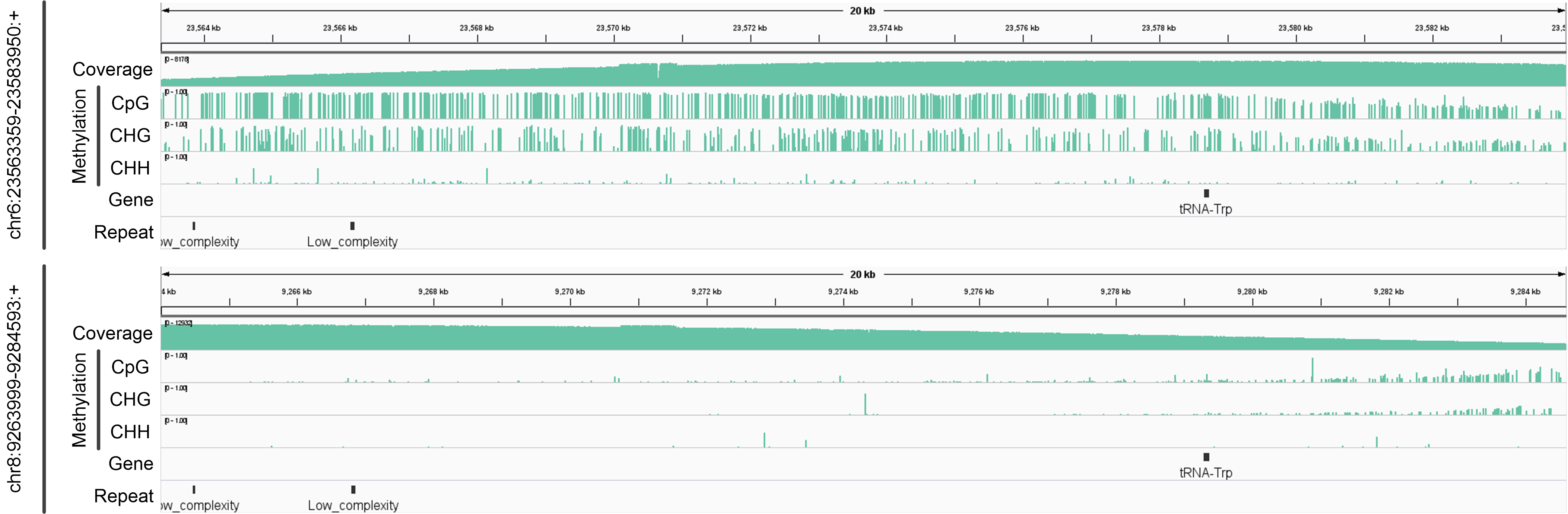

**Supplementary Fig. 27** Genome browser view of a differentially methylated repeat pair (chr6:23563359-23583950: +, chr8: 9263999-9284593:+) in *O. sativa* (sample2).

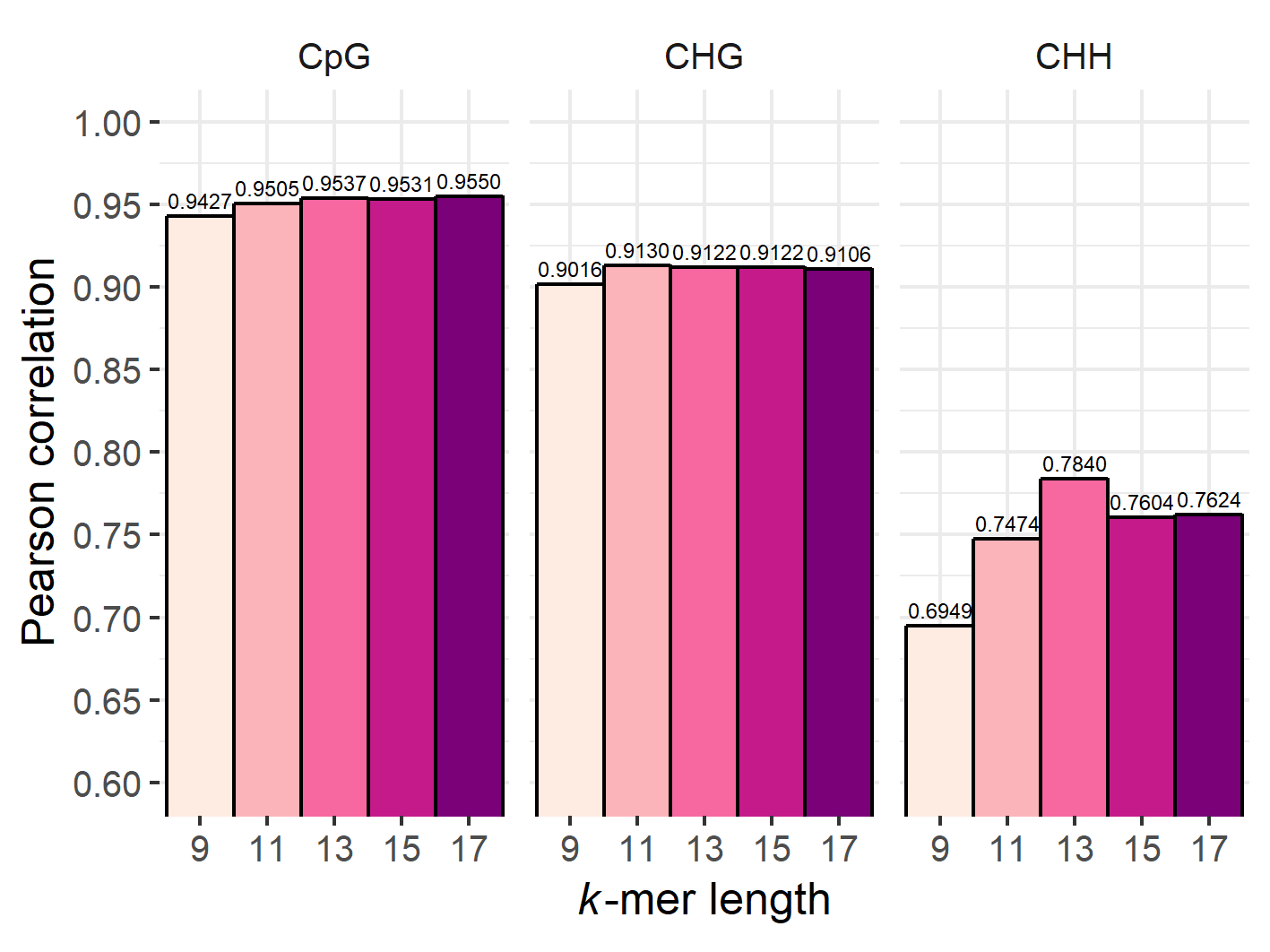

**Supplementary Fig. 28** *k*-mer length tuning of DeepSignal-plant. The training samples are extracted from ~500× Nanopore reads of *A. thaliana*. Pearson correlations are calculated using the results from ~20× Nanopore reads and three bisulfite replicates of *A. thaliana*.

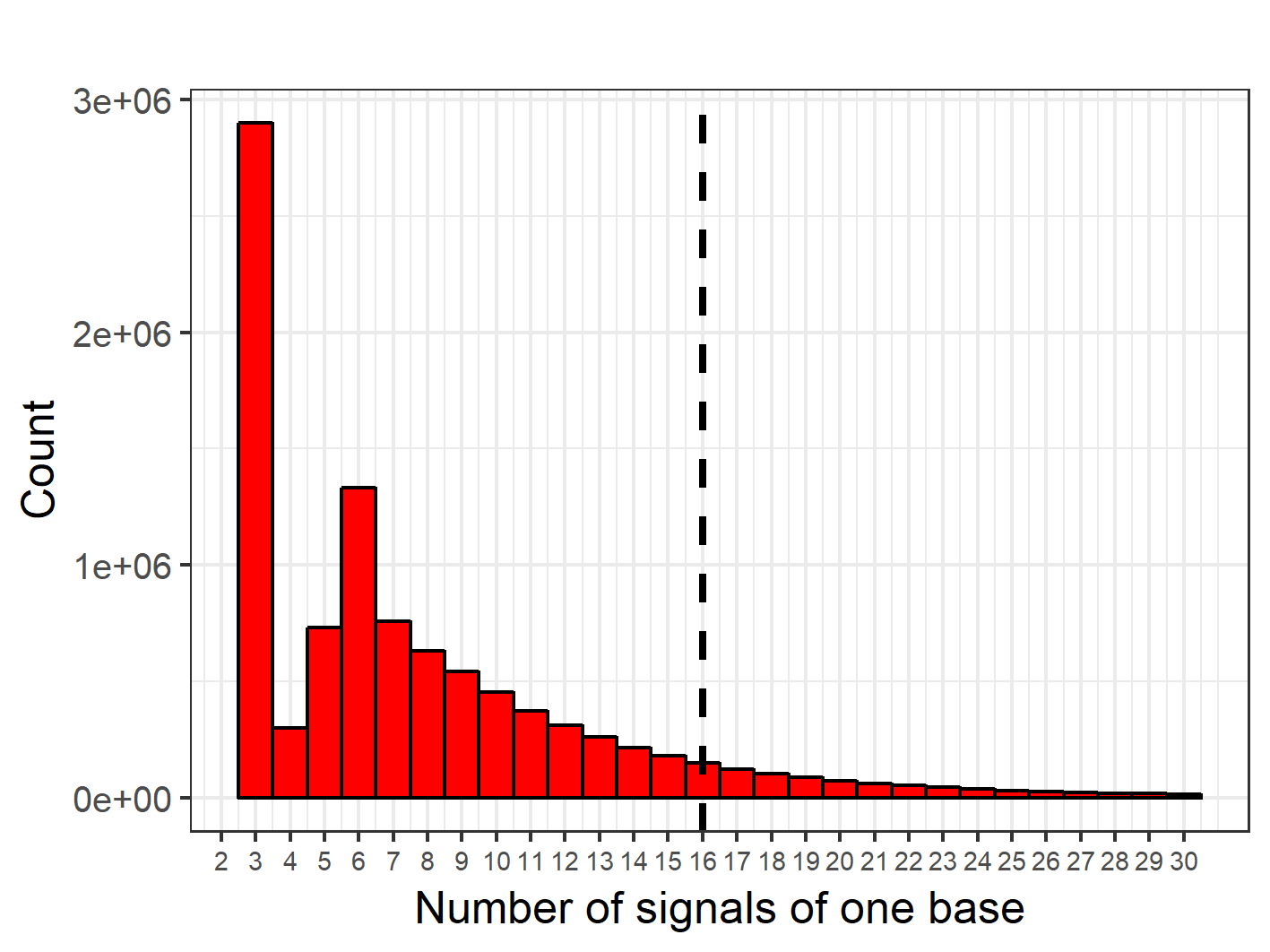

**Supplementary Fig. 29** Number of signals of 10 million randomly selected bases. Suppose *u* and *σ* are mean and standard deviation of the number of signals, the dash line indicates approximately *u+σ* signals.

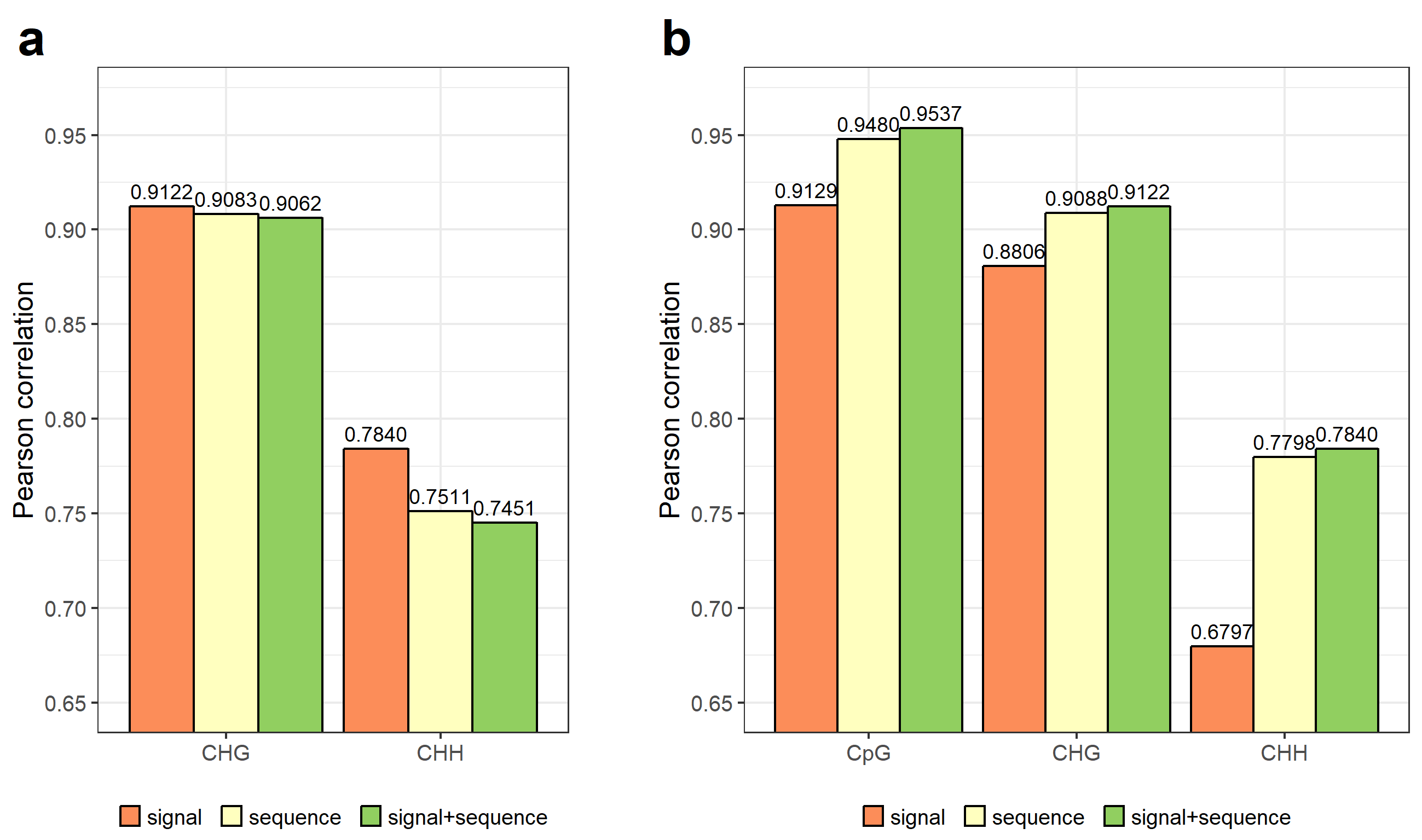

**Supplementary Fig. 30** Feature selection of DeepSignal-plant to denoise training samples and call methylation (The training samples are extracted from ~500× Nanopore reads. Pearson correlations are calculated using the results from ~20× Nanopore reads and three bisulfite replicates of *A. thaliana*.). **a:** Evaluation of different features to denoise training samples (After denoising training samples, the training samples are used to training models for calling methylation by using signal+sequence features). **b:** Evaluation of different features to call methylation (All the training samples are balanced first and then denoised (for CHG and CHH motif only) by using only signal features).

#### Supplementary Tables

**Supplementary Table 1.** Number of CpG, CHG, CHH sites which bisulfite sequencing (Bismark) and Nanopore sequencing (DeepSignal-plant) can detected in *A. thaliana* and *O. sativa*. We count the sites from 5 chromosomes of *A. thaliana* genome and 12 chromosomes of *O. sativa* genome. Sites from both forward and complement strand of the genomes are counted. In bisulfite sequencing of *A. thaliana*, we count sites which satisfy *cov*>=1 or 5 in at least 1 replicate. In Nanopore sequencing, we count sites which satisfy *cov*>=1 or 5 from ~100x tested reads. *cov*=coverage.

| species | motif | genome |  | bisulfite | |  | Nanopore | |
| --- | --- | --- | --- | --- | --- | --- | --- | --- |
|  |  |  |  | *cov*>=1 | *cov*>=5 |  | *cov*>=1 | *cov*>=5 |
| *A. thaliana* | CpG | 5,567,714 |  | 5,487,342 | 5,468,996 |  | 5,549,652 | 5,521,044 |
|  | CHG | 6,093,647 |  | 6,014,437 | 5,996,330 |  | 6,079,079 | 6,063,014 |
|  | CHH | 31,198,155 |  | 30,774,058 | 30,653,400 |  | 31,106,922 | 31,011,252 |
| *O. sativa* (rep1) | CpG | 30,817,376 |  | 29,594,582 | 28,712,658 |  | 30,714,046 | 30,498,978 |
|  | CHG | 27,376,461 |  | 26,418,299 | 25,767,463 |  | 27,316,646 | 27,196,935 |
|  | CHH | 104,355,374 |  | 100,637,175 | 97,252,858 |  | 104,123,055 | 103,686,499 |
| *O. sativa* (rep2) | CpG | 30,817,376 |  | 29,755,811 | 28,542,701 |  | 30,711,556 | 30,486,515 |
|  | CHG | 27,376,461 |  | 26,562,610 | 25,722,292 |  | 27,315,000 | 27,195,722 |
|  | CHH | 104,355,374 |  | 101,378,360 | 98,481,274 |  | 104,117,684 | 103,686,107 |

**Supplementary Table 2.** Number of parameters in the model architecture of DeepSignal and DeepSignal-plant.

| feature source | DeepSignal | |  | DeepSignal-plant | |
| --- | --- | --- | --- | --- | --- |
|  | architecture | No. of parameters |  | architecture | No. of parameters |
| sequence features | 3-layer BiLSTM | 3,026,944 |  | 1-layer BiLSTM +  1 fully connected layer | 173,248 |
| signal features | 11 inception blocks | 991,680 |  | 1-layer BiLSTM +  1 fully connected layer | 182,400 |
| concatenated | 2 fully connected layers | 36,397,088 |  | 3-layer BiLSTM +  2 fully connected layers | 4,338,434 |
| - | total | 40,415,712 |  | total | 4,694,082 |

**Supplementary Table 3.** Number of high-confidence sites in *A. thaliana* (3 technical replicates) and *O. sativa* (2 biological replicates). (A site is considered to be methylated with high confidence if the site is covered with at least 5 reads and has at least 0.9 methylation frequency. A site is considered to be unmethylated with high confidence if it has at least five mapped reads and the methylation frequency is 0. Numbers in bold indicate the number of sites we select to train models.)

| motif | state | *A. thaliana* | | | | |  | *O. sativa* | |
| --- | --- | --- | --- | --- | --- | --- | --- | --- | --- |
|  |  | rep1 | rep2 | rep3 | intersection | union |  | rep1 | rep2 |
| CpG | methylated | 546,320 | 543,200 | 553,574 | **233,528** | 882,728 |  | **13,367,759** | 13,405,189 |
|  | unmethylated | 3,120,040 | 3,019,453 | 3,111,175 | **2,257,533** | 3,630,296 |  | **9,380,609** | 10,053,549 |
| CHG | methylated | 35,512 | 34,258 | 36,253 | 12,482 | **65,076** |  | **2,845,705** | 2,833,756 |
|  | unmethylated | 4,161,281 | 4,018,022 | 4,149,921 | **3,018,418** | 4,823,546 |  | **10,213,164** | 11,523,142 |
| CHH | methylated | 7,577 | 6,962 | 7,722 | 1,226 | **16,434** |  | **148,789** | 131,885 |
|  | unmethylated | 22,285,718 | 21,400,918 | 22,212,460 | **15,717,929** | 26,382,803 |  | **53,916,702** | 63,447,979 |

**Supplementary Table 4.** Comparison of the number of unique *k*-mers in high-confidence methylated and unmethylated sites for training (*k*=13).

| motif | *A. thaliana* | | |  | | *O. sativa* | | |
| --- | --- | --- | --- | --- | --- | --- | --- | --- |
|  | methylated | unmethylated | intersection |  | methylated | | unmethylated | intersection |
| CpG | 179,687 | 1,343,098 | 88,496 |  | 2,635,394 | | 2,808,801 | 1,905,205 |
| CHG | 43,713 | 1,386,150 | 29,856 |  | 736,876 | | 2,503,090 | 637,048 |
| CHH | 13,107 | 5,294,855 | 10,354 |  | 65,033 | | 8,340,086 | 62,066 |

**Supplementary Table 5.** Comparison of per-site methylation frequencies predicted by DeepSignal-plant and Megalodon from Nanopore sequencing with the results calculated from bisulfite sequencing in *A. thaliana* and *O. sativa*. ~100× Nanopore reads of *A. thaliana*, *O. sativa* (rep1), and *O. sativa* (rep2) were used, respectively. Models of DeepSignal-plant and Megalodon were trained by using combined reads of *A. thaliana* and *O. sativa*. *r*: Pearson correlation; *r^2^*: coefficient of determination; *ρ*: Spearman correlation; *RMSE*: root mean square error.

| species | motif | method | *r* | *r^2^* | *ρ* | *RMSE* |
| --- | --- | --- | --- | --- | --- | --- |
| *A. thaliana* | CpG | DeepSignal-plant | 0.9840 | 0.9683 | 0.8144 | 0.0699 |
|  |  | Megalodon | 0.9654 | 0.9321 | 0.8186 | 0.1101 |
|  | CHG | DeepSignal-plant | 0.9638 | 0.9289 | 0.6907 | 0.0567 |
|  |  | Megalodon | 0.9353 | 0.8749 | 0.7876 | 0.0819 |
|  | CHH | DeepSignal-plant | 0.8975 | 0.8054 | 0.5391 | 0.0466 |
|  |  | Megalodon | 0.5783 | 0.3344 | 0.4302 | 0.0904 |
| *O. sativa* (rep1) | CpG | DeepSignal-plant | 0.9919 | 0.9838 | 0.8496 | 0.0623 |
|  |  | Megalodon | 0.9871 | 0.9744 | 0.8453 | 0.0770 |
|  | CHG | DeepSignal-plant | 0.9603 | 0.9222 | 0.8765 | 0.1043 |
|  |  | Megalodon | 0.9551 | 0.9121 | 0.9035 | 0.1185 |
|  | CHH | DeepSignal-plant | 0.8563 | 0.7332 | 0.5124 | 0.0644 |
|  |  | Megalodon | 0.7157 | 0.5122 | 0.4156 | 0.0890 |
| *O. sativa* (rep2) | CpG | DeepSignal-plant | 0.9918 | 0.9837 | 0.8597 | 0.0617 |
|  |  | Megalodon | 0.9814 | 0.9632 | 0.8584 | 0.0952 |
|  | CHG | DeepSignal-plant | 0.9603 | 0.9221 | 0.8888 | 0.1042 |
|  |  | Megalodon | 0.9498 | 0.9021 | 0.9098 | 0.1214 |
|  | CHH | DeepSignal-plant | 0.8686 | 0.7545 | 0.5431 | 0.0582 |
|  |  | Megalodon | 0.7220 | 0.5212 | 0.4382 | 0.0766 |

**Supplementary Table 6.** Comparison of per-site methylation frequencies predicted by DeepSignal-plant and Megalodon from Nanopore sequencing with the results calculated from bisulfite sequencing in *B. nigra*. ~78× Nanopore reads were used. Models of DeepSignal-plant and Megalodon were trained by using combined reads of *A. thaliana* and *O. sativa*. *r*: Pearson correlation; *r^2^*: coefficient of determination; *ρ*: Spearman correlation; *RMSE*: root mean square error.

| motif | method | *r* | *r^2^* | *ρ* | *RMSE* |
| --- | --- | --- | --- | --- | --- |
| CpG | DeepSignal-plant | 0.9693 | 0.9396 | 0.6832 | 0.1022 |
|  | Megalodon | 0.9637 | 0.9287 | 0.6442 | 0.1070 |
| CHG | DeepSignal-plant | 0.8941 | 0.7994 | 0.8645 | 0.1390 |
|  | Megalodon | 0.8944 | 0.7999 | 0.8691 | 0.1416 |
| CHH | DeepSignal-plant | 0.7488 | 0.5607 | 0.5421 | 0.1024 |
|  | Megalodon | 0.5307 | 0.2817 | 0.3924 | 0.1379 |

**Supplementary Table 7.** Number of CpG, CHG, CHH sites detected by bisulfite sequencing (~20×, BSMAP) and Nanopore sequencing (~78×, DeepSignal-plant) in *B. nigra*. Sites from both forward and complement strand of 8 chromosomes of *B. nigra* are counted. *cov*=coverage.

| motif | genome |  | bisulfite | |  | Nanopore | |
| --- | --- | --- | --- | --- | --- | --- | --- |
|  |  |  | *cov*>=1 | *cov*>=5 |  | *cov*>=1 | *cov*>=5 |
| CpG | 27,476,356 |  | 21,231,907 | 15,485,216 |  | 27,346,210 | 26,936,082 |
| CHG | 28,459,137 |  | 21,744,279 | 15,551,935 |  | 28,424,635 | 28,293,960 |
| CHH | 133018020 |  | 102,908,660 | 65,592,026 |  | 132,917,689 | 132,603,400 |

**Supplementary Table 8.** Number of repeat pairs in *A. thaliana* and *O. sativa*. Differentially methylated repeat pairs are based on the results of DeepSignal-plant. “total” counts repeat pairs which contain at least 1 cytosine in the corresponding motif. “differential” counts repeat pairs which contain at least 10% differentially methylated cytosines in the corresponding motif.

| species | motif | repeat pairs | | | | | | | |
| --- | --- | --- | --- | --- | --- | --- | --- | --- | --- |
|  |  | length>=100 | |  | length>=1000 | |  | length>=10000 | |
|  |  | total | differential |  | total | differential |  | total | differential |
| A. thaliana | C | 1,104 | 101 |  | 356 | 9 |  | 46 | 0 |
|  | CpG | 936 | 97 |  | 356 | 28 |  | 46 | 0 |
|  | CHG | 938 | 78 |  | 356 | 13 |  | 46 | 0 |
|  | CHH | 1,103 | 57 |  | 356 | 2 |  | 46 | 0 |
| O. sativa (rep1) | C | 26,508 | 1,528 |  | 10,358 | 241 |  | 256 | 6 |
|  | CpG | 24,964 | 1,199 |  | 10,358 | 354 |  | 256 | 19 |
|  | CHG | 24,941 | 2,449 |  | 10,358 | 458 |  | 256 | 10 |
|  | CHH | 26,476 | 873 |  | 10,358 | 34 |  | 256 | 0 |
| O. sativa (rep2) | C | 26,508 | 1,640 |  | 10,358 | 250 |  | 256 | 5 |
|  | CpG | 24,964 | 1,235 |  | 10,358 | 357 |  | 256 | 16 |
|  | CHG | 24,941 | 2,514 |  | 10,358 | 479 |  | 256 | 10 |
|  | CHH | 26,476 | 935 |  | 10,358 | 32 |  | 256 | 0 |

#### Supplementary Notes

**Supplementary Note 1:** **Model architecture of DeepSignal-plant**

(1) A bidirectional LSTM layer

A bidirectional LSTM layer includes a forward LSTM and a backward LSTM to catch both forward and reverse flow of features. Let *x_1_, x_2_,…, x_t_* are a sequence of features. For sequence features used in DeepSignal-plant, each time step $x_{i}$ contains four features: the nucleotide base, the mean, standard deviation and the number of mapped signals of the base. For signal features in DeepSignal-plant, each time step $x_{i}$ contains *m* features which are *m* signals of the current base. A LSTM will recursively calculate the hidden layer *h* as follows:

$i_{t}=sigmoid(W_{xi}x_{t}+{W_{hi}h}_{t-1}+{W_{ci}⨀c}_{t-1}+b_{i})$	(1)

$f_{t}=sigmoid(W_{xf}x_{t}+{W_{hf}h}_{t-1}+{W_{cf}⨀c}_{t-1}+b_{f})$	(2)

$c_{t}=f_{t}{\odot c}_{t-1}+i_{t}\odot tanh(W_{xc}x_{t}+{W_{hc}⨀c}_{t}+b_{c})$	(3)

$o_{t}=sigmoid(W_{xo}x_{t}+{W_{ho}h}_{t-1}+{W_{co}⨀c}_{t}+b_{o})$	(4)

$h_{t}=o_{t}\odot tanh(c_{t})$	(5)

where *W* and *b* are weights and biases in the model. *x* is the input vector; *i* is the activation vector of input gate; *f* is the activation vector of forget gate; *c* is the cell state vector; *o* is the activation vector of output gate; and *h* is the output vector of the LSTM hidden unit. Current output $h_{t}$of LSTM hidden unit depends on the input $x_{t},$ previous state $h_{t-1}$, and previous information stored in cell. Then, the outputs of forward and backward LSTM are combined:

$z_{t}=h_{t,F}\bigoplus h_{t,B}$	(6)

(2) Softmax activation function

In DeepSignal-plant, softmax activation function is used to predict the methylated and unmethylated probabilities of one sample as follows:

$softmax\left( x_{i} \right)=\frac{e^{x_{i}}}{\sum_{j=0}^{1} e^{x_{j}}}, i=0 or 1$	(7)

where *x_0_* and *x_1_* are two outputs from the former fully connected layer.

(3) The cross-entropy loss function used during training is as follows:

$L=z*-log\left( y \right)+\left( 1-z \right)*-log(1-y)$	(8)

where *z* is the true label vector and *y* is the predicted probability vector output from softmax function.

**Supplementary Note 2: Hyperparameters and feature selection of DeepSignal-plant**

In DeepSignal-plant, we use bidirectional long short-term memory (BiLSTM) layers with 256 hidden size and tanh activation function. We use one BiLSTM layer to receive sequence features and signal features, respectively. Three BiLSTM layers are used to process the concatenated features. By using the randomly selected ~500× *A. thaliana* reads for training and another ~20× *A. thaliana* reads for testing, we test the performances of different *k*-mer lengths (Supplementary Fig. 28). According to the results, we set *k*=13 as default. By testing the number of signals of 10 million bases randomly selected from reads of *A. thaliana*, we set *m* (*i.e.*, number of signals of each base) to 16, which is great than the number of signals of most bases (Supplementary Fig. 29).

During training, we use dropout probability of 0.5 at each dropout layer. And we use a batch size of 512 and an initial learning rate 0.001. The learning rate is adopted by Adam optimizer and decayed by a factor of 0.1 after each two epochs. The parameter *betas* in Adam optimizer is set to (0.9, 0.999) as default. We train at least 5 epochs and at most 10 epochs at each training process.

DeepSignal-plant uses two groups of features (sequence features and signal features) to predict methylation state of one targeted site. Besides test different *k*-mer lengths, we further use the reads of *A. thaliana* to test the effectiveness of the two groups of features in denoising training samples and calling methylation. As shown in Supplementary Fig. 30a, for CHG and CHH motif, using signal features to denoise training samples gets the best performances. For calling methylation of all three motifs, using signal features gets the worst performance and using both features gets the highest performance (Supplementary Fig. 30b).
